# An ancient transcription factor functions as a master regulator of primary cilia formation

**DOI:** 10.1101/2025.11.28.691250

**Authors:** Weihua Wang, Xiqi Zhang, Yaxuan Qiu, Xiangrui Meng, Sitong Cheng, Yutong Chen, Siqi Liu, Wenhui Chen, Jiayan Yi, Xiwen You, Hongni Liu, Junqiao Xing, Cheng Xu, Haochen Jiang, Haibo Wang, Guangmei Tian, Zhangfeng Hu

## Abstract

How ancient transcription factors are repurposed during evolution to drive the functional diversification of conserved organelles remains a fundamental question in biology. Although highly conserved in basic morphology, eukaryotic cilia vary extensively in their sizes and functions. Previously, we showed that an evolutionarily ancient transcription factor, X chromosome-associated protein 5 (Xap5), controls motile ciliogenic transcriptional programs during mouse spermatogenesis (Wang et al., 2025). Here we show that Xap5 functions as a master regulator of primary ciliogenesis. In NIH/3T3 fibroblasts, Xap5 associates with the nuclear protein Non-POU domain-containing octamer-binding protein (Nono) to form a regulatory module that promotes Sox5 and Sox9 expression. Genetic ablation of *Xap5* or *Nono* shortens primary cilia, and loss of Sox5 disrupts the downstream ciliogenic program, consistent with the established role of Sox9. In mIMCD-3 renal epithelial cells, Xap5 loss reduces both ciliation frequency and cilium length, whereas Sox5 and Sox9 respond in the opposite direction. Collectively, our results provide new insight into how an ancient transcription factor is repurposed through shared and context-specific downstream programs, and motivate testing whether altered Xap5-dependent gene regulation contributes to human ciliopathies.

## Introduction

A central challenge in evolutionary biology is to understand how a limited number of ancient conserved genes generate tremendous cellular diversity through functional redeployment^1-3^. This challenge is particularly evident in the roles of master transcription factors, which must be precisely regulated across different cellular contexts to control specific developmental programs^4-7^. Such functional plasticity is thought to arise through evolutionary co-option, the process by which existing regulators are recruited into new regulatory modules to drive lineage- or organelle-specific innovations^8-10^. Elucidating the molecular strategies underlying this repurposing is essential for understanding how complex biological systems evolve, and begins with defining how individual conserved regulators operate in different cellular contexts. The biogenesis of the cilium—an evolutionarily ancient organelle that plays key roles in signaling and motility^11-14^—provides an important model for investigating this fundamental question, and its dysfunction can lead to a class of genetic disorders known as ciliopathies^15^.

The assembly of the cilium, or ciliogenesis, requires the precise transcriptional coordination of hundreds of genes^16-18^. In metazoans, the regulatory factor X (Rfx) family and the transcription factor Forkhead box protein J1 (Foxj1) have been identified as key regulators of ciliogenesis^19-23^. Rfx factors regulate the expression of genes essential for building the core ciliary structure^19,20^, and are required for the formation of sensory and primary cilia in *C. elegans*^24^, *Drosophila*^25^ and the developing mouse nervous system^26^, while Foxj1 directs the motile ciliogenic program^27,28^. Upstream of these core factors, the regulatory hierarchy is best characterized for the motile cilia of multiciliated cells^21,22^. How the upstream, cell-type-specific transcriptional programs are organized, particularly for the non-motile primary cilium found on most somatic cells, remains poorly understood^12,29^.

The conserved transcription factor Xap5 serves as a prime example of this challenge. Our previous work showed that Xap5 regulates the assembly of motile appendages—dynamic structures that generate movement by beating in fluid environments—in the unicellular alga *Chlamydomonas*^30^, and controls ciliary transcriptional programs during mouse spermatogenesis^31^. These findings established Xap5’s role in motile ciliogenic programs^32^, yet Xap5 is also broadly expressed across mouse tissues^31^, posing a biological enigma: why is a regulator of motile ciliogenesis expressed so broadly? It remains to be determined whether Xap5 plays a role in primary ciliogenesis in somatic cells and, if so, through which downstream effectors it acts.

In this study, we specifically address these questions by characterizing the function of Xap5 in primary ciliogenesis using NIH/3T3 mouse fibroblasts, a standard model for primary cilium formation upon serum starvation^33^, and mIMCD-3 renal epithelial cells. Here, we identify the Non-POU domain-containing octamer-binding protein (Nono) as a critical interacting partner of Xap5. Our integrated analyses demonstrate that Xap5 and Nono form a regulatory module that promotes Sox5 and Sox9 expression and is required for normal cilium length in NIH/3T3 cells. While Sox9 has been previously implicated in primary ciliogenesis^34-36^, our findings identify Sox5 as a previously unrecognized regulator of primary cilium length. In mIMCD-3 cells, Xap5 loss also impaired primary ciliogenesis, with transcriptional responses that were partly shared with and partly distinct from those in NIH/3T3 cells. Together, these findings identify Xap5 as a master regulator of primary cilia formation. Moreover, they provide a framework for how ancient transcription factors are functionally repurposed across cellular contexts.

## Results

### Xap5 is required for normal primary cilium length in NIH/3T3 cells

To investigate the role of Xap5 in primary cilia formation, we first determined its subcellular localization in NIH/3T3 cells. Immunofluorescence staining with a specific anti-Xap5 antibody revealed that endogenous Xap5 is predominantly nuclear (Fig. 1a), consistent with our previous work. We confirmed this finding via ectopic expression of a Flag-tagged Xap5 construct (Fig. 1b). Consistent with its nuclear localization, bioinformatic analysis predicted a putative nuclear localization signal (NLS) within the Xap5 protein (Supplementary Fig. 1a). To examine the contribution of this sequence to nuclear localization, we generated an NLS mutant, designated ΔNLS (Supplementary Fig. 1a). The mutant showed pronounced cytoplasmic localization, indicating that disruption of the NLS impaired the normal nuclear localization of Xap5 (Fig. 1b).

**Fig. 1.**
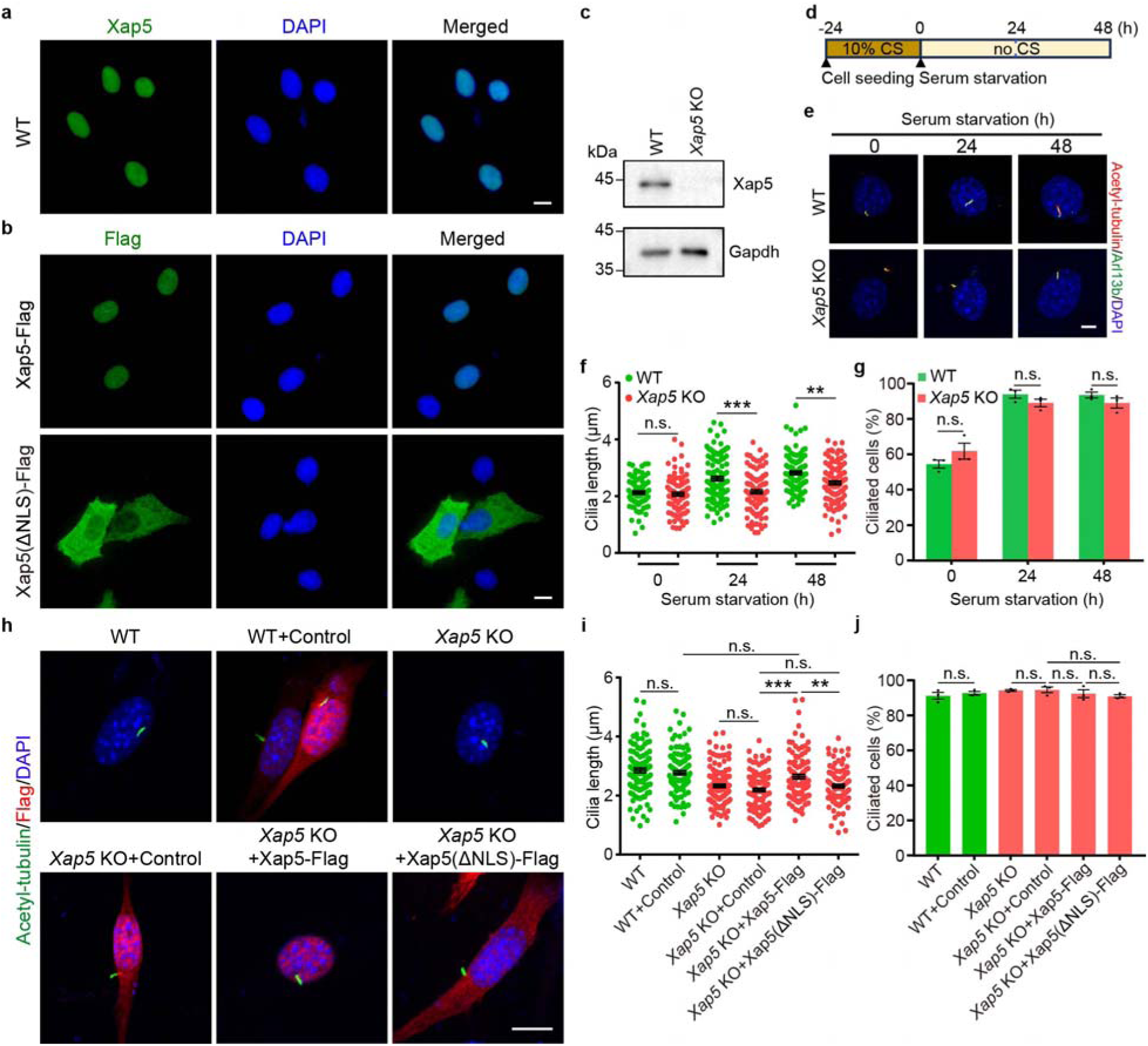
Xap5 is a nuclear protein required for normal primary cilium length in NIH/3T3 cells. **a** Immunofluorescence analysis of endogenous Xap5 (green) in NIH/3T3 cells. Nuclei were counterstained with DAPI (blue). Scale bar, 10 μm. **b** Validation of the Xap5 nuclear localization signal (NLS). Representative images of NIH/3T3 cells expressing either Flag-tagged wild-type Xap5 (WT) or an NLS mutant, designated ΔNLS. Flag-tagged proteins are shown in green and DAPI-stained nuclei in blue. Scale bar, 10 μm. **c** Western blot analysis confirming the loss of detectable Xap5 protein in the *Xap5* knockout (KO) clonal cell line. Gapdh serves as a loading control. The blot shown is representative of three independent biological replicates. **d** Schematic illustrating the time-course assay for primary cilia assembly induced by serum starvation. CS, calf serum. **e** Representative immunofluorescence images of cilia from WT and *Xap5* KO cells at the indicated time points. Cilia were co-stained for acetylated tubulin (red) and Arl13b (green). Nuclei were counterstained with DAPI (blue). Scale bar, 5 μm. **f** Quantification of primary cilia length in WT and *Xap5* KO cells over a 48-h time course of serum starvation. Data are presented as means ± SEM (n = 3 independent experiments). **g** Quantification of the percentage of ciliated cells as in **f**. Data are presented as mean ± SEM (n = 3 independent experiments). **h** Representative immunofluorescence images of WT and *Xap5* KO NIH/3T3 cells under the indicated expression conditions following 24 h of serum starvation. Control denotes the GFP-Flag expression control. Cells were stained for acetylated tubulin (green) and Flag-tagged proteins (red), with nuclei counterstained with DAPI (blue). Scale bar, 10 μm. **i** Quantification of primary cilium length in the groups shown in **h**. Data are presented as mean ± SEM (n = 3 independent experiments). **j** Quantification of the percentage of ciliated cells as in **i**. Data are presented as mean ± SEM (n = 3 independent experiments). *P* values were determined by multiple two-tailed unpaired t-tests. n.s., not significant; \*\**P* < 0.01; \*\*\**P* < 0.001. Source data are provided as a Source Data file.

To investigate the function of Xap5, we generated *Xap5* knockout (KO) NIH/3T3 cell lines using the CRISPR/Cas9 system^37^ (Supplementary Fig. 1b). Western blotting showed loss of detectable Xap5 protein (Fig. 1c). Upon inducing ciliogenesis via serum starvation (Fig. 1d), we first observed a striking defect in ciliary length (Fig. 1e). Although comparable to wild-type (WT) cells before starvation, *Xap5* KO cells exhibited a significant shortening of their primary cilia post-starvation (Fig. 1e, f). To determine if this length defect stemmed from an overall failure in ciliogenesis, we next quantified the ciliation frequency. Interestingly, both WT and *Xap5* KO cells responded robustly to the starvation stimulus, showing a comparable increase in the percentage of ciliated cells over 48 h (Fig. 1g). Thus, under the conditions examined in NIH/3T3 cells, Xap5 loss reduced ciliary length without a detectable change in ciliation frequency.

To assess the functional contribution of the Xap5 NLS, we expressed Xap5-Flag or Xap5(ΔNLS)-Flag in *Xap5* KO NIH/3T3 cells. GFP-Flag served as an expression control in WT and KO cells, and cilia were assessed after 24 h of serum starvation (Fig. 1h). No significant differences in cilium length were detected between the GFP-Flag groups and their respective WT or KO controls. In KO cells, Xap5-Flag increased cilium length relative to GFP-Flag. Cilium length did not differ significantly between the KO + Xap5-Flag and WT + GFP-Flag groups. The KO + Xap5(ΔNLS)-Flag group had shorter cilia than the KO + Xap5-Flag group. Its cilium length did not differ significantly from that of the KO + GFP-Flag group (Fig. 1i). No significant differences in ciliation frequency were detected in the indicated comparisons (Fig. 1j). Together with the localization data, these findings support a contribution of nuclear localization to cilium-length regulation by Xap5, although other effects of the substitutions remain possible.

### Xap5 forms a nuclear regulatory module with its cofactor Nono

Given that nuclear localization contributes to cilium-length regulation by Xap5 (Fig. 1i), we next searched for candidate Xap5-associated proteins by immunoprecipitating endogenous Xap5 from NIH/3T3 cells and analyzing the immunoprecipitate by mass spectrometry (IP-MS). The analysis yielded 92 protein identifications, including Xap5 itself, Nono, Sfpq and core histones (Supplementary Data 1). Nono and Sfpq belong to the *Drosophila* behavior/human splicing (DBHS) family and can form heterodimers^38^. Because Xap5 has been implicated in RNA polymerase II recruitment in *Chlamydomonas*^30^, we prioritized Nono, which has been reported to support RNA polymerase II recruitment to cAMP-responsive promoters^39^.

As a prerequisite for their interaction, we first confirmed that Nono is also a predominantly nuclear protein in NIH/3T3 cells, a localization we validated for both the endogenous protein via immunofluorescence and an ectopically expressed Nono-GFP fusion protein (Fig. 2a). To biochemically validate their physical association, we next performed endogenous co- immunoprecipitation (Co-IP) assays. These results confirmed a robust and specific interaction, as immunoprecipitation of Xap5 specifically co-precipitated Nono (Fig. 2b). Taken together, these localization and co-immunoprecipitation data provide strong evidence that Xap5 and Nono form a bona fide regulatory complex within the nucleus.

**Fig. 2.**
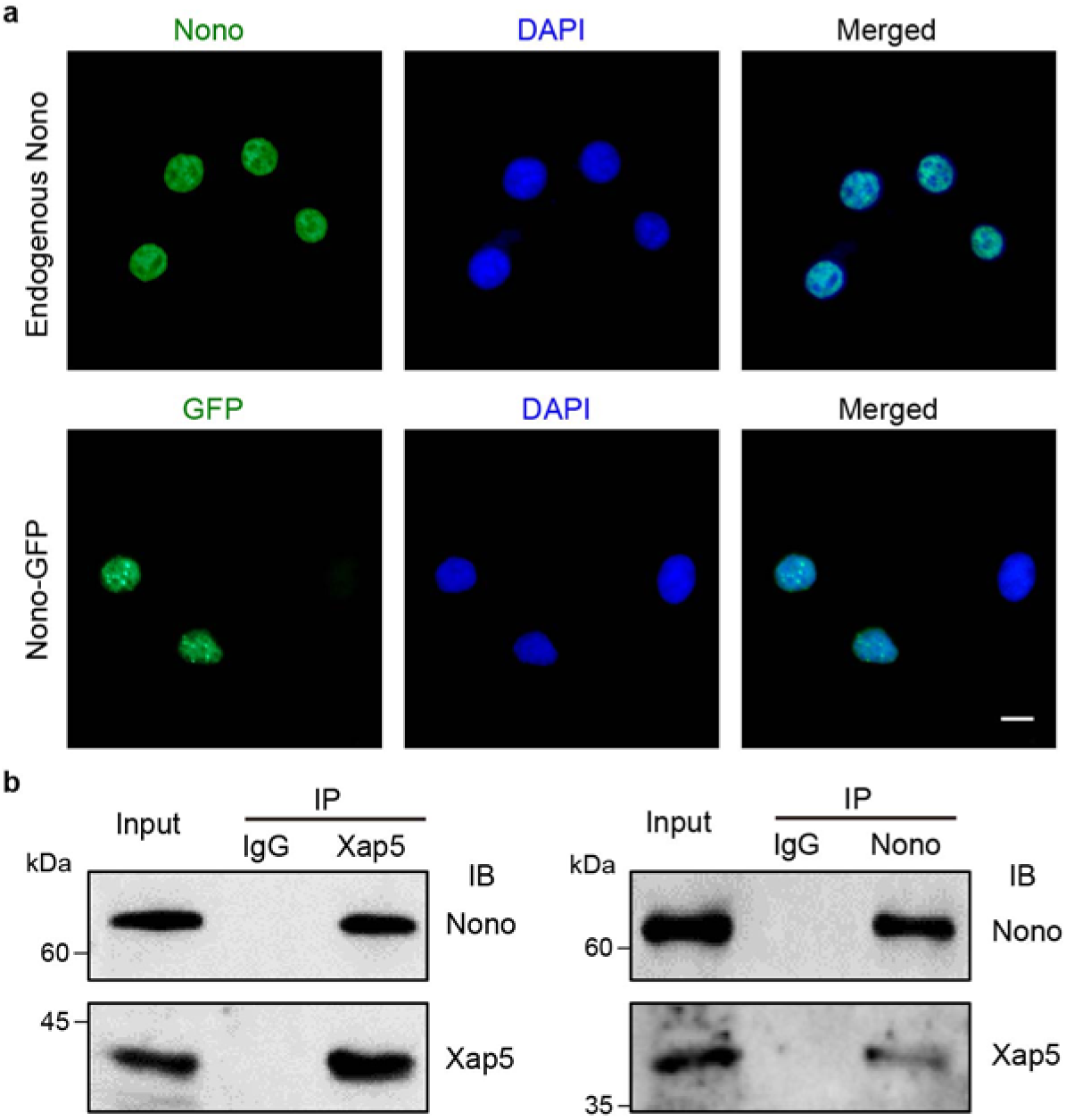
Xap5 forms a nuclear complex with its cofactor Nono. **a** Representative immunofluorescence images showing the predominantly nuclear localization of Nono in NIH/3T3 cells. Top: Endogenous Nono detected with a specific anti-Nono antibody (green). Bottom: Ectopically expressed Nono-GFP fusion protein (green). Nuclei were counterstained with DAPI (blue). Scale bar, 10 μm. **b** Endogenous co-immunoprecipitation (Co-IP) of Xap5 and Nono from NIH/3T3 cell lysates. Left: Immunoprecipitation (IP) with an anti-Xap5 antibody specifically co-precipitated endogenous Nono. Right: Conversely, IP with an anti-Nono antibody co-precipitated endogenous Xap5. Control IgG was used as a negative control for the IP. The resulting precipitates (IP) and input lysates were analyzed by western blotting (IB) with the indicated antibodies. Data are representative of three independent experiments. Source data are provided as a Source Data file.

### The Xap5–Nono module is required for normal primary cilium length

Given that Nono forms a complex with Xap5, we asked whether loss of Nono produces the same ciliary phenotype as loss of Xap5. To test this, we generated *Nono* KO NIH/3T3 cells using the CRISPR/Cas9 system (Supplementary Fig. 2a), and immunoblotting showed loss of detectable Nono protein (Fig. 3a). Indeed, upon serum starvation, *Nono* KO cells phenocopied the *Xap5* KO. They exhibited significantly shorter primary cilia compared to wild-type (WT) controls, yet displayed a comparable increase in ciliation frequency (Fig. 3b-d). Notably, in addition to the ciliary defects, we observed a significant increase in nuclear area in *Nono* KO cells (Fig. 3b; Supplementary Fig. 2b). Together, these findings demonstrate that Nono, like Xap5, is required for normal cilium length, whereas its loss did not detectably change ciliation frequency under the conditions examined. Nono may also have a broader role in maintaining nuclear architecture.

**Fig. 3.**
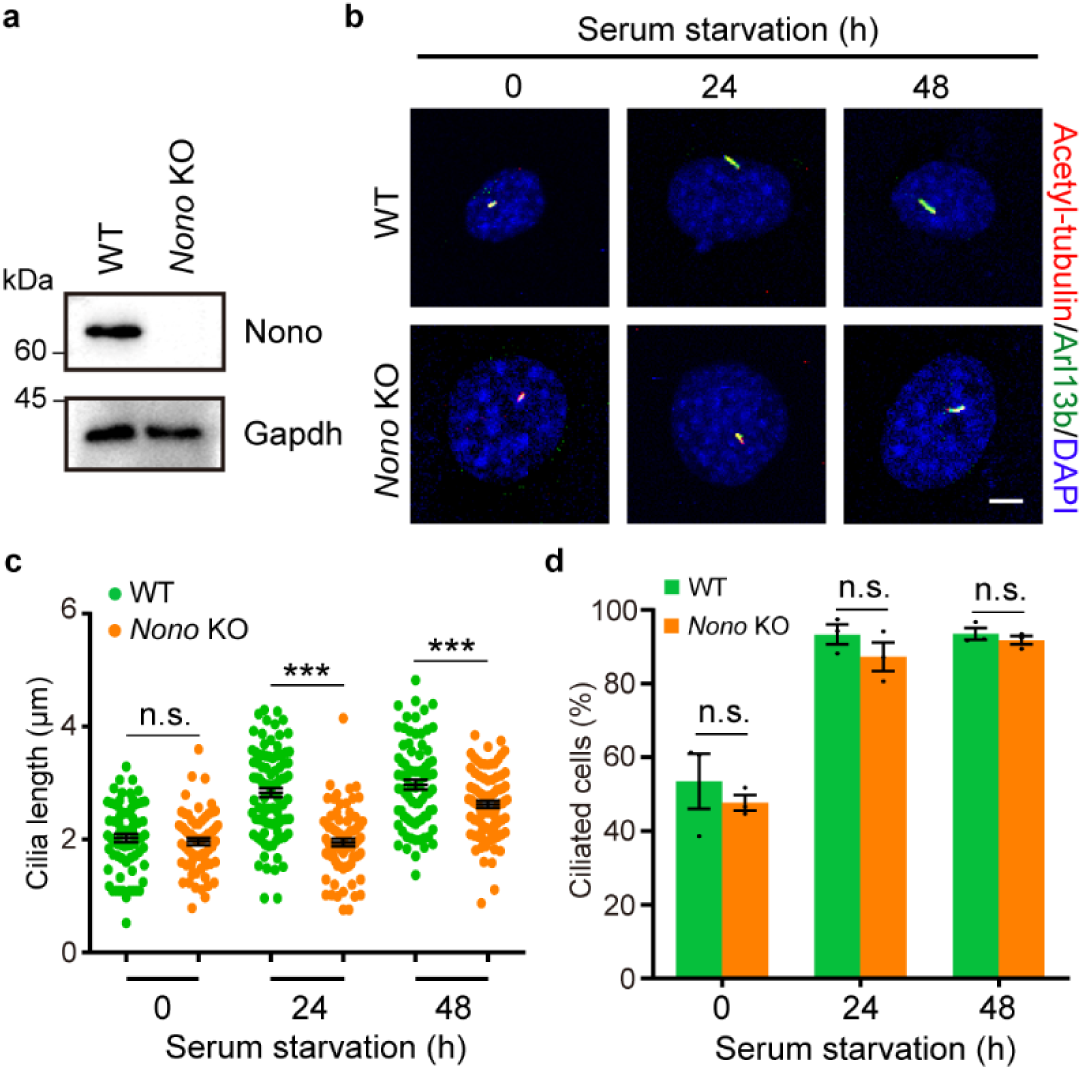
Loss of Nono phenocopies the cilium-length defect caused by Xap5 loss. **a** Western blot analysis confirming the loss of detectable Nono protein in *Nono* KO cell lysates compared to WT. The blot shown is representative of three independent biological replicates. Gapdh serves as a loading control. **b** Representative immunofluorescence images of primary cilia in WT and *Nono* KO cells at the indicated time points. Cilia were co-stained for acetyl-tubulin (red) and Arl13b (green). Nuclei were counterstained with DAPI (blue). Scale bar, 5 μm. **c** Quantification of primary cilia length in WT and *Nono* KO cells over a 48-h time course of serum starvation. Data are presented as mean ± SEM (n = 3 independent experiments). **d** Quantification of the percentage of ciliated cells as in **c**. Data are presented as mean ± SEM (n = 3 independent experiments). All statistical significance between WT and KO at each time point was determined by multiple two-tailed unpaired t-tests. n.s., not significant; \*\**P* < 0.01; \*\*\**P* < 0.001. Source data are provided as a Source Data file.

### The Xap5–Nono module co-regulates a downstream ciliogenic transcriptional cascade

To determine if the Xap5–Nono module cooperates at the transcriptional level, we performed RNA-sequencing (RNA-seq) on WT, *Xap5* KO, and *Nono* KO NIH/3T3 cells following 24 h of serum starvation, a time point where the phenotype was robustly established. This analysis revealed widespread transcriptional changes; *Xap5* KO cells showed 1,116 upregulated and 1,071 downregulated genes, while *Nono* KO cells had 1,424 upregulated and 1,043 downregulated genes relative to WT (fold change ≥ 1.5; Supplementary Fig. 3a, b; Supplementary Data 2, 3). Gene Ontology (GO) analyses performed separately on the genes downregulated in each KO identified cilium-related and transcription-related categories among the enriched terms (Fig. 4a, b; Supplementary Fig. 3c, d).

**Fig. 4.**
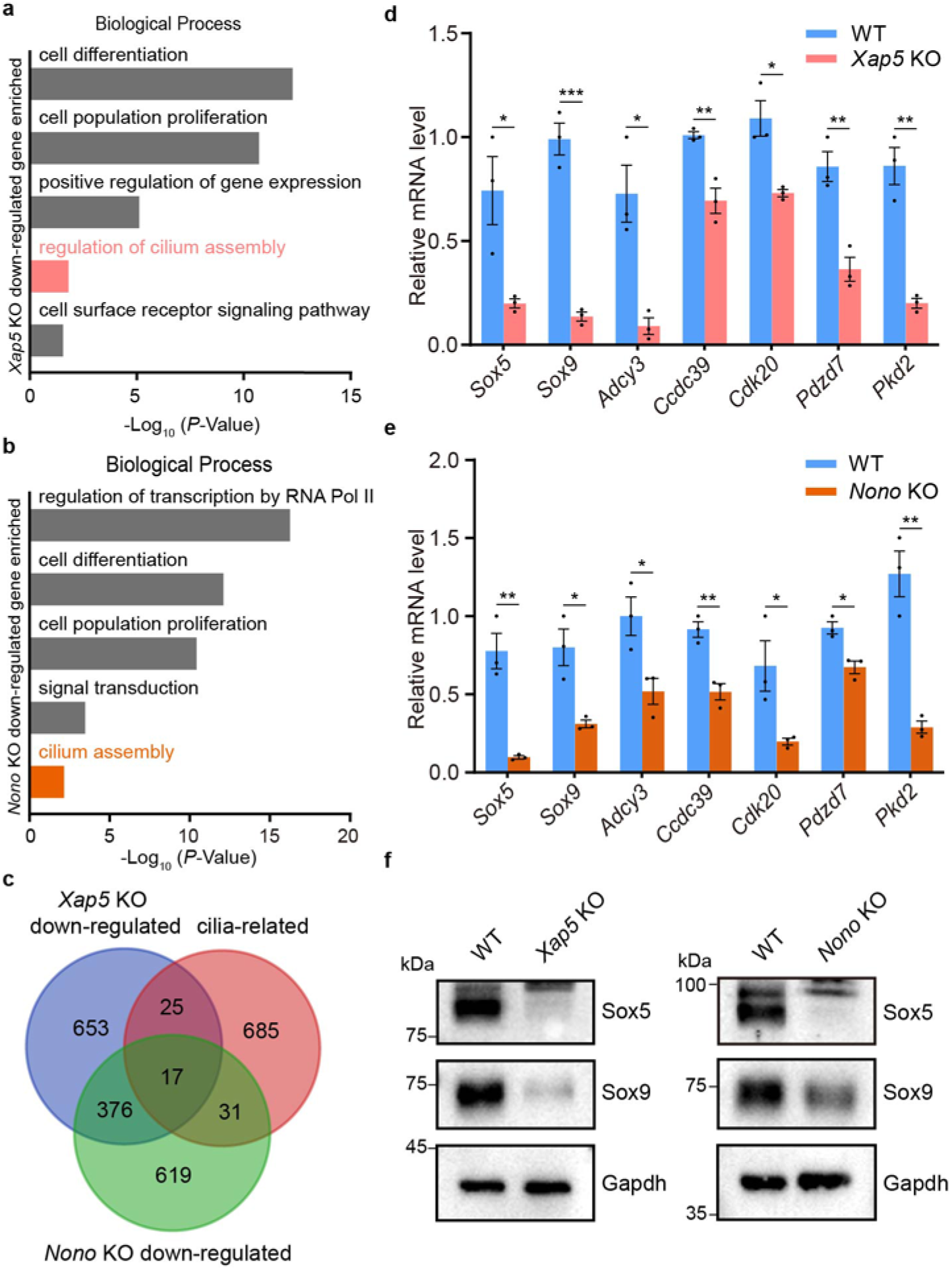
Xap5 and Nono co-regulate a ciliogenic transcriptional network. All analyses were performed on NIH/3T3 cells following 24 h of serum starvation. **a, b** Representative enriched Gene Ontology (GO) terms (Biological Process) for genes downregulated in *Xap5* KO (**a**) and *Nono* KO (**b**) cells. **c** Venn diagram illustrating the overlap between genes downregulated in *Xap5* KO cells, *Nono* KO cells, and a curated list of cilia-associated genes (Supplementary Data 4). **d, e** qPCR validation of shared, downregulated ciliary target genes in *Xap5* KO (**d**) and *Nono* KO (**e**) cells. Data are presented as mean ± SEM from three independent experiments. *P* values were determined by two-tailed unpaired t-test. \**P* < 0.05, \*\**P* < 0.01, \*\*\**P* < 0.001. **f** Representative western blots showing reduced protein levels of Sox5 and Sox9 in *Xap5* KO (left) and *Nono* KO (right) cells compared to their respective WT controls. Gapdh serves as a loading control. Blots are representative of three independent experiments. Source data are provided as a Source Data file.

We previously showed that Xap5 regulates Foxj1 and Rfx2 during early spermatogenesis^31^, but neither gene was among the differentially expressed genes in *Xap5* KO NIH/3T3 cells (Supplementary Data 2). We examined their expression by RT-qPCR, together with the ciliogenic transcription factor gene *Rfx3* and the IFT-A component gene *Ift122*. *Rfx2* mRNA levels were significantly higher in *Xap5* KO cells than in WT cells, whereas *Foxj1* mRNA levels did not differ significantly between genotypes. By contrast, *Rfx3* and *Ift122* mRNA levels were significantly lower (Supplementary Fig. 3e). *Rfx3* was also among the genes downregulated in *Nono* KO cells in the RNA-seq analysis (Supplementary Data 3). Immunoblotting showed that Foxj1 and Rfx2 are expressed in NIH/3T3 cells and that Rfx2 levels were not reduced in *Xap5* KO cells (Supplementary Fig. 3f). Thus, unlike during early spermatogenesis^31^, Xap5 loss in NIH/3T3 cells did not lower *Foxj1* or *Rfx2* mRNA levels but lowered those of other ciliogenic genes, such as *Rfx3* and *Ift122*.

*Nono* mRNA levels were modestly but significantly higher in *Xap5* KO cells, and Nono protein signal was not reduced (Supplementary Fig. 3e, f). Thus, Xap5 is not required for Nono expression. Given that loss of either Xap5 or Nono shortens primary cilia, we compared the downregulated gene set against a comprehensive ciliary gene list (Supplementary Data 4), which was assembled from the SYSCILIA gold standard (SCGSv2)^18^ and manual literature curation. This comparison confirmed a significant overlap between the two gene sets (Fig. 4c). We validated several of these shared targets—including the transcription factors *Sox5* and *Sox9*, and the ciliary component *Pkd2*—by qPCR (Fig. 4d, e).

Among this cohort of downregulated genes, two transcription factors, Sox5 and Sox9, stood out. Sox9 is a known regulator of primary cilia in other contexts^34-36^, and both factors are known to cooperate in developmental programs^40^. This made them candidates for further investigation as potential downstream effectors in our system. Consistent with this hypothesis, western blot analysis confirmed that the protein levels of both Sox5 and Sox9 were markedly reduced in *Xap5* and *Nono* KO NIH/3T3 cells (Fig. 4f). These data strongly indicate that Xap5 and Nono co-regulate a transcriptional network that includes the downstream effectors Sox5 and Sox9, thereby driving primary cilia formation.

### Xap5 and Nono occupy the Sox5 and Sox9 promoter regions in NIH/3T3 cells

To determine whether Xap5 and Nono occupy regulatory regions of the genes identified above, we profiled each protein separately by CUT&Tag^41^ in NIH/3T3 cells after 24 h of serum starvation (Supplementary Data 5, 6). Signal for both proteins was enriched around transcription start sites (TSSs). Approximately 30% of Xap5 peaks and 50% of Nono peaks were annotated as promoter-proximal (Fig. 5a; Supplementary Fig. 4a-d). Genes assigned to peaks included both upregulated and downregulated genes in the corresponding KO (Supplementary Fig. 4e, f).

**Fig. 5.**
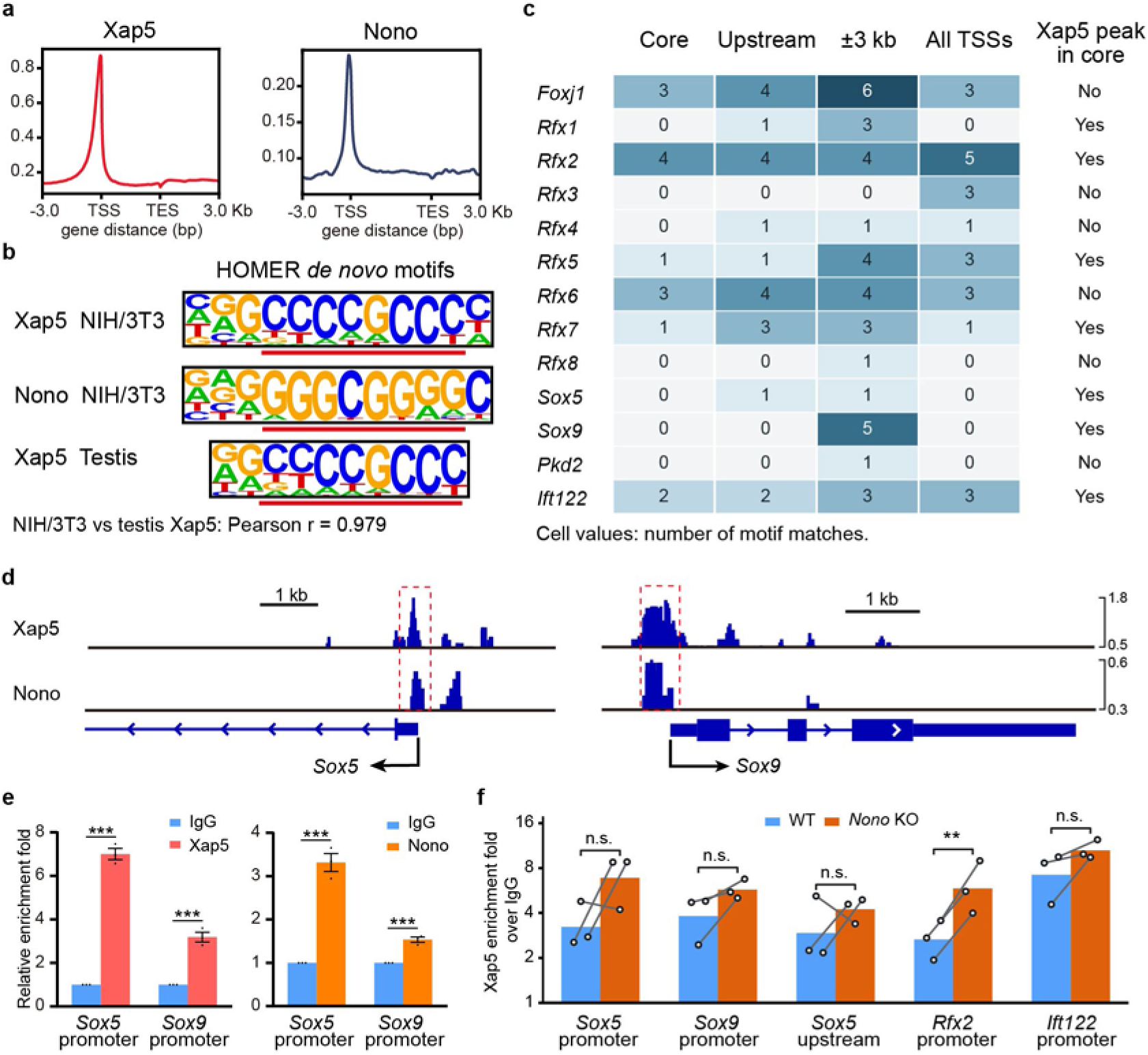
Xap5 and Nono co-occupy the regulatory regions of ciliary genes. **a** Average CUT&Tag profiles of Xap5 and Nono across gene bodies and 3-kb flanking regions. TSS, transcription start site; TES, transcription end site. **b** *De novo* motifs from Xap5 and Nono CUT&Tag peaks in NIH/3T3 cells and the previously reported Xap5 motif from mouse testis^31^. **c** Counts of Xap5-associated motif matches in four promoter windows: Core, −500 to +250 bp relative to the representative TSS; Upstream, −3 kb to the TSS; ±3 kb; and All TSSs, the union of core windows around all annotated TSSs. The final column indicates Xap5 peak overlap with the representative core promoter. **d** IGV tracks of Xap5 and Nono at the *Sox5* and *Sox9* loci. Red dashed boxes highlight promoter-proximal enrichment. Scale bars, 1 kb. **e** CUT&Tag–qPCR enrichment of Xap5 (left) and Nono (right) at the *Sox5* and *Sox9* promoters, relative to IgG set to 1. **f** Xap5 CUT&Tag–qPCR enrichment over matched IgG in WT and *Nono* KO cells at the indicated regions, plotted on a log_2_ scale. Lines connect paired samples. Bars show mean ± SEM (**e**) or geometric means (**f**) from three independent experiments. Panels **e** and **f** used separate WT samples. P values were determined using two-tailed Student’s t-tests: unpaired on linear enrichment values (**e**) and paired on log_2_ enrichment (**f**), without multiple-comparison correction. n.s., not significant; \*\**P* < 0.01; \*\*\**P* < 0.001. Source data are provided as a Source Data file.

*De novo* motif analysis identified GC-rich motifs in both peak sets (Fig. 5b). The Xap5-associated motifs from NIH/3T3 cells and mouse testis^31^ each ranked first among 880 candidate motifs in reciprocal TOMTOM searches and aligned across all ten positions of the shorter motif (Pearson r = 0.979). Of 45,495 eligible Xap5 peak windows from WT NIH/3T3 cells, 37.95% contained a motif match, compared with 20.38% and 24.32% in whole-window and locally binned dinucleotide-preserving shuffles, respectively. Among windows containing a match, the highest-scoring sites were more often near the peak summit than in either shuffled reference (central match weight, 22.44% versus 10.09% and 13.11%; Supplementary Fig. 5a-c). Thus, Xap5 peaks in NIH/3T3 cells and testis share a similar GC-rich motif, which in NIH/3T3 peaks is enriched and concentrated near summits relative to both dinucleotide-preserving shuffled references.

We next scanned the promoters of *Foxj1* and all eight Rfx genes with the NIH/3T3 motif (Supplementary Fig. 5d). The representative core promoters of *Foxj1*, *Rfx2*, *Rfx5*, *Rfx6* and *Rfx7* contained matches; the other four Rfx genes, including *Rfx3*, had matches only in wider windows or around alternative TSSs (Fig. 5c). Among genes assigned to a WT Xap5 peak and differentially expressed in *Xap5* KO cells (Supplementary Fig. 4e), 590 of 926 evaluable downregulated genes and 554 of 849 evaluable upregulated genes had a core-promoter match (Supplementary Data 5). For example, *Ift122* was downregulated in *Xap5* KO cells (Supplementary Data 2), and its core promoter contained both a motif match and an Xap5 peak (Fig. 5c); the *Rfx2* core promoter had both features, whereas *Rfx2* mRNA was higher in *Xap5* KO cells by RT–qPCR (Supplementary Fig. 3e). In a model of 7,068 genes (expression groups from a separate RNA-seq reanalysis), neither the downregulated nor the upregulated group differed significantly from the stable group in the odds of a core-promoter match after adjustment for GC content, CpG density and WT expression (odds ratios 0.86 and 1.10; Holm-adjusted P = 0.121 and 0.269, respectively; Supplementary Data 5). Thus, core-promoter matches were common among Xap5 peak-assigned genes whose expression changed in either direction, but motif presence was not significantly associated with expression-response category in the adjusted model.

Of the 282 genes assigned to peaks for both proteins and downregulated in both KO lines, 14 were on the curated list of 758 cilia-associated genes, including Sox5 (Supplementary Figs. 4e, f and 5e; Supplementary Data 4). Sox9, also downregulated in both KO lines (Fig. 4d–f), was outside this intersection because its promoter-proximal Nono peak was not assigned to Sox9 in the peak-to-gene annotation (Supplementary Data 6). At the loci of both candidate downstream effectors, Sox5 and Sox9, the Xap5 and Nono profiles showed promoter-proximal signal, and CUT&Tag–qPCR detected enrichment of each protein over IgG (Fig. 5d, e). Xap5 and Nono therefore each occupy the *Sox5* and *Sox9* promoters.

We next asked whether the lower Sox5 and Sox9 expression in *Nono* KO cells was accompanied by lower Xap5 enrichment. In three experiments, each including both genotypes, we assayed five regions: the *Sox5* and *Sox9* promoter regions shown in Fig. 5e; a region in a separate Xap5 peak upstream of *Sox5* (Supplementary Fig. 5f); and the *Rfx2* and *Ift122* promoters, where the core windows each contain motif matches and overlap an Xap5 peak (Fig. 5c). Of the five amplicons, only the *Sox5* upstream amplicon contained a complete motif match at the scan threshold. *Xap5* mRNA did not differ significantly between genotypes, and Xap5 protein was detected in both (Supplementary Fig. 5g, h). In both genotypes, anti-Xap5 signal exceeded matched IgG at every region in every experiment. KO/WT ratios of geometric mean enrichment ranged from 1.44 to 2.20. Enrichment relative to IgG was significantly higher at the *Rfx2* promoter, whereas the anti-Xap5 signal alone did not differ significantly between genotypes (Fig. 5f). Thus, Sox5 and Sox9 expression declined after Nono loss while Xap5 remained enriched at their assayed regions. Together, these results identify *Sox5* and *Sox9* as shared targets of Xap5 and Nono in NIH/3T3 cells.

### Sox5 acts downstream of the Xap5–Nono module to control primary cilium length

Our finding that Xap5 and Nono co-regulate Sox5 and Sox9 prompted us to investigate their respective roles in ciliogenesis. Given the established role of Sox9 in primary ciliogenesis^34-36^, we focused our functional analysis on the uncharacterized role of Sox5. To this end, we generated *Sox5* KO NIH/3T3 cells using the CRISPR/Cas9 system (Fig. 6a; Supplementary Fig. 6).

**Fig. 6.**
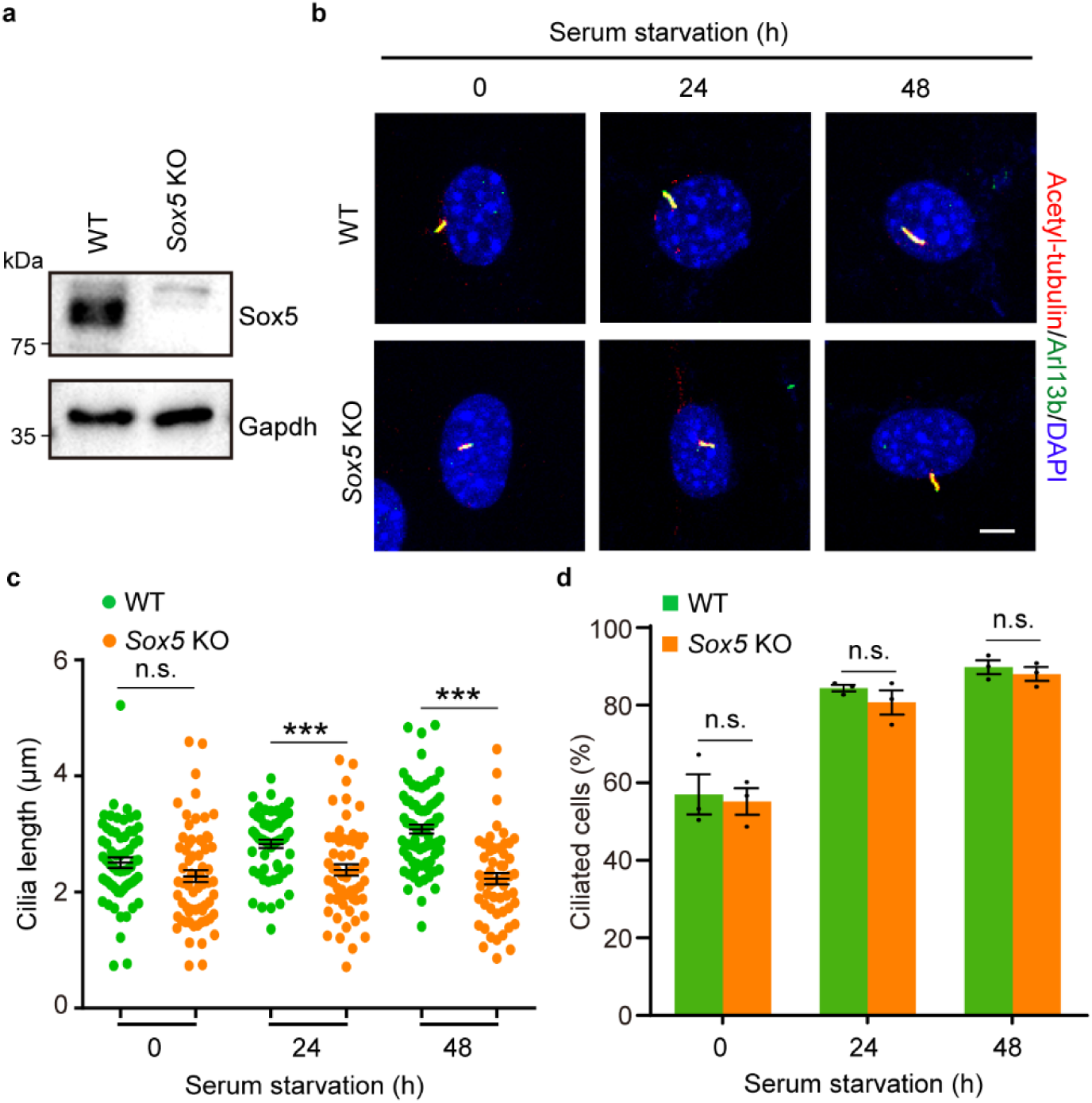
Downstream effector Sox5 is required for normal primary cilium length. **a** Western blot analysis of Sox5 in WT and *Sox5* KO cell lysates. The blot shown is representative of three independent biological replicates. Gapdh serves as a loading control. **b** Representative immunofluorescence images of primary cilia in WT and *Sox5* KO cells after serum starvation. Cilia were co-stained for acetylated tubulin (red) and Arl13b (green). Nuclei were counterstained with DAPI (blue). Scale bar, 5 μm. **c** Quantification of primary cilia length in WT and *Sox5* KO cells over a 48-h time course of serum starvation. Data are presented as mean ± SEM (n = 3 independent experiments). **d** Quantification of the percentage of ciliated cells as in **c**. Data are presented as mean ± SEM from three independent experiments. *P* values were determined by two-tailed unpaired t-test. n.s., not significant; \*\*\**P* < 0.001. Source data are provided as a Source Data file.

Analysis after serum starvation revealed that *Sox5* KO cells phenocopied the upstream knockouts. *Sox5* KO cells exhibited significantly shorter primary cilia than WT controls (Fig. 6b, c), while both genotypes showed a comparable increase in ciliation frequency (Fig. 6d). These results identify Sox5 as a downstream effector required for normal primary cilium length. Together with the established role of Sox9 in primary ciliogenesis^34-36^, these data support a model in which the Xap5–Nono module acts through the downstream transcription factors Sox5 and Sox9 to promote normal primary cilia.

### Sox5 executes a shared ciliogenic program downstream of the Xap5–Nono complex

To confirm that Sox5 functions transcriptionally downstream of the Xap5–Nono complex, we performed RNA-seq on *Sox5* KO and WT cells after 24 h of serum starvation. The analysis identified 2,124 upregulated and 2,105 downregulated genes in *Sox5* KO cells (fold change ≥ 1.5; Supplementary Fig. 7a; Supplementary Data 7). To assess the global impact on ciliary pathways, we first performed Gene Set Enrichment Analysis (GSEA)^42^ on the ranked gene list from the *Sox5* KO versus WT comparison, which revealed a significant enrichment of the “Cilium organization” pathway (Fig. 7a). Consistent with this, GO analysis of the strictly defined downregulated gene set (fold change ≥ 1.5) also highlighted terms related to cilium assembly (Fig. 7b; Supplementary Fig. 7b). Furthermore, direct comparison showed an overlap between the full set of genes dysregulated by Sox5 loss and our curated ciliary gene list (Supplementary Fig. 7c), reinforcing the strong link between Sox5 and ciliogenesis.

**Fig. 7.**
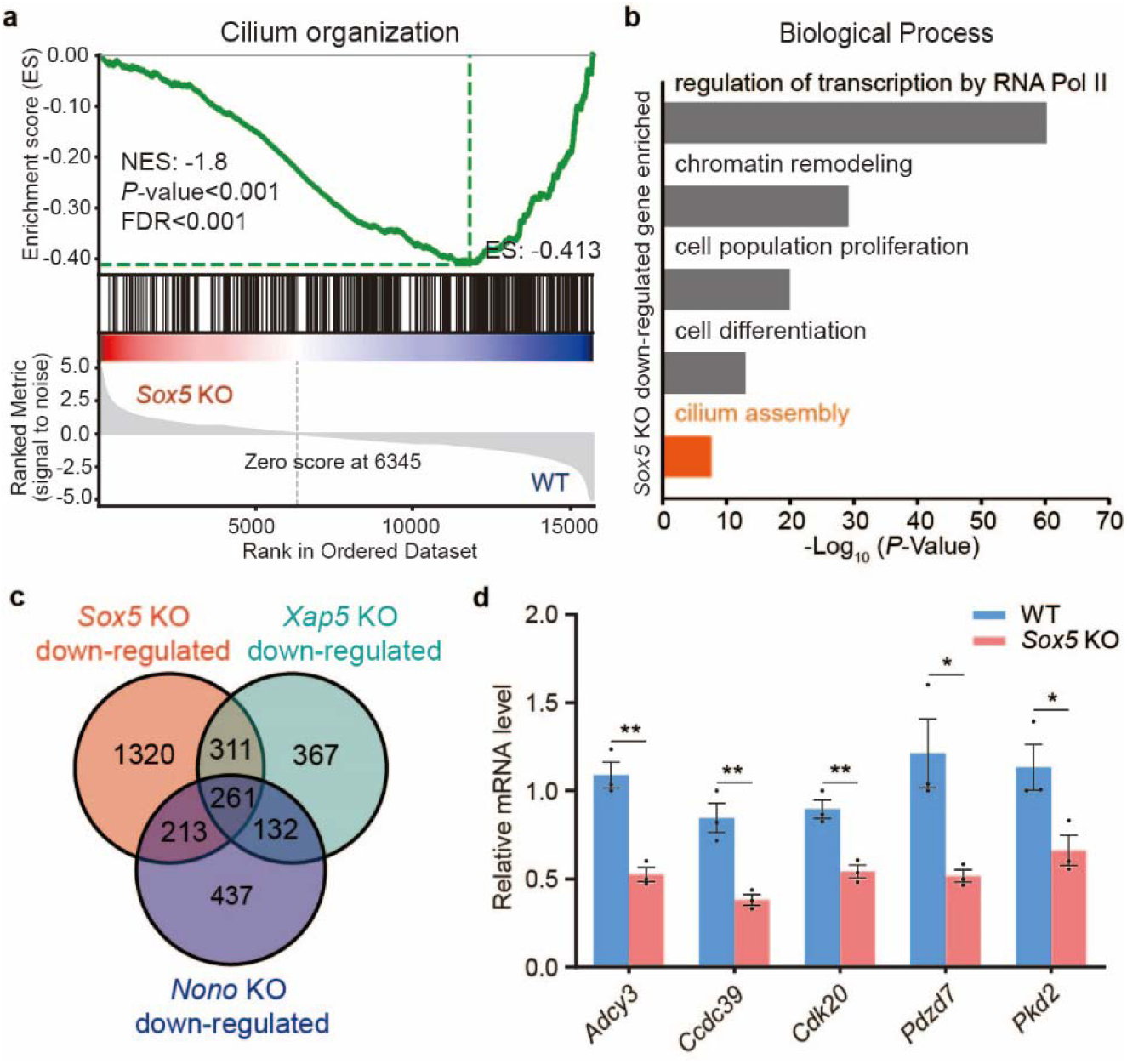
Sox5 executes a ciliogenic transcriptional program downstream of the Xap5–Nono complex. **a** Gene Set Enrichment Analysis (GSEA) plot for the “cilium organization” pathway, performed on the ranked gene list from the *Sox5* KO versus WT comparison. **b** Representative enriched GO terms (Biological Process) for genes significantly downregulated (fold change ≥ 1.5) in *Sox5* KO cells. **c** Venn diagram illustrating the overlap among genes downregulated in *Xap5* KO, *Nono* KO, and *Sox5* KO cells, identifying a common set of co-regulated genes. **d** qPCR validation confirming the reduced expression of shared downstream target genes in *Sox5* KO cells compared to WT. Data are presented as mean ± SEM from three independent experiments. *P* values were determined by two-tailed unpaired t-test. \**P* < 0.05, \*\**P* < 0.01. Source data are provided as a Source Data file.

Critically, to place Sox5 within the Xap5–Nono pathway, we compared the downregulated gene sets across all three KO lines. A Venn diagram revealed a substantial overlap among them (Fig. 7c). We confirmed this shared transcriptional dependency by qPCR, validating that several key ciliary genes—including *Adcy3*, *Ccdc39*, and *Pkd2*—were indeed downregulated across all three KO models (Figs. 4d, e and 7d). These results provide strong evidence that Sox5 acts as a key transcriptional mediator downstream of Xap5 and Nono, executing a common program of ciliary gene expression required for normal cilium length in NIH/3T3 cells.

### Xap5 loss impairs primary ciliogenesis in mIMCD-3 renal epithelial cells

To determine whether the contribution of Xap5 to primary ciliogenesis extends beyond fibroblasts, we examined *Xap5* KO mIMCD-3 mouse renal epithelial cells. Sanger sequencing of the targeted region identified a 2-bp deletion predicted to introduce a premature stop codon in the *Xap5* KO clone (Supplementary Fig. 8). Immunoblotting showed loss of detectable Xap5 protein (Fig. 8a). Compared with WT controls, *Xap5* KO cells exhibited significantly shorter primary cilia both before serum starvation and after 24 h of starvation (Fig. 8b, c). The percentage of ciliated cells was also significantly reduced at both time points (Fig. 8d). Thus, in this epithelial model, Xap5 loss reduced both cilium length and ciliation frequency.

**Fig. 8.**
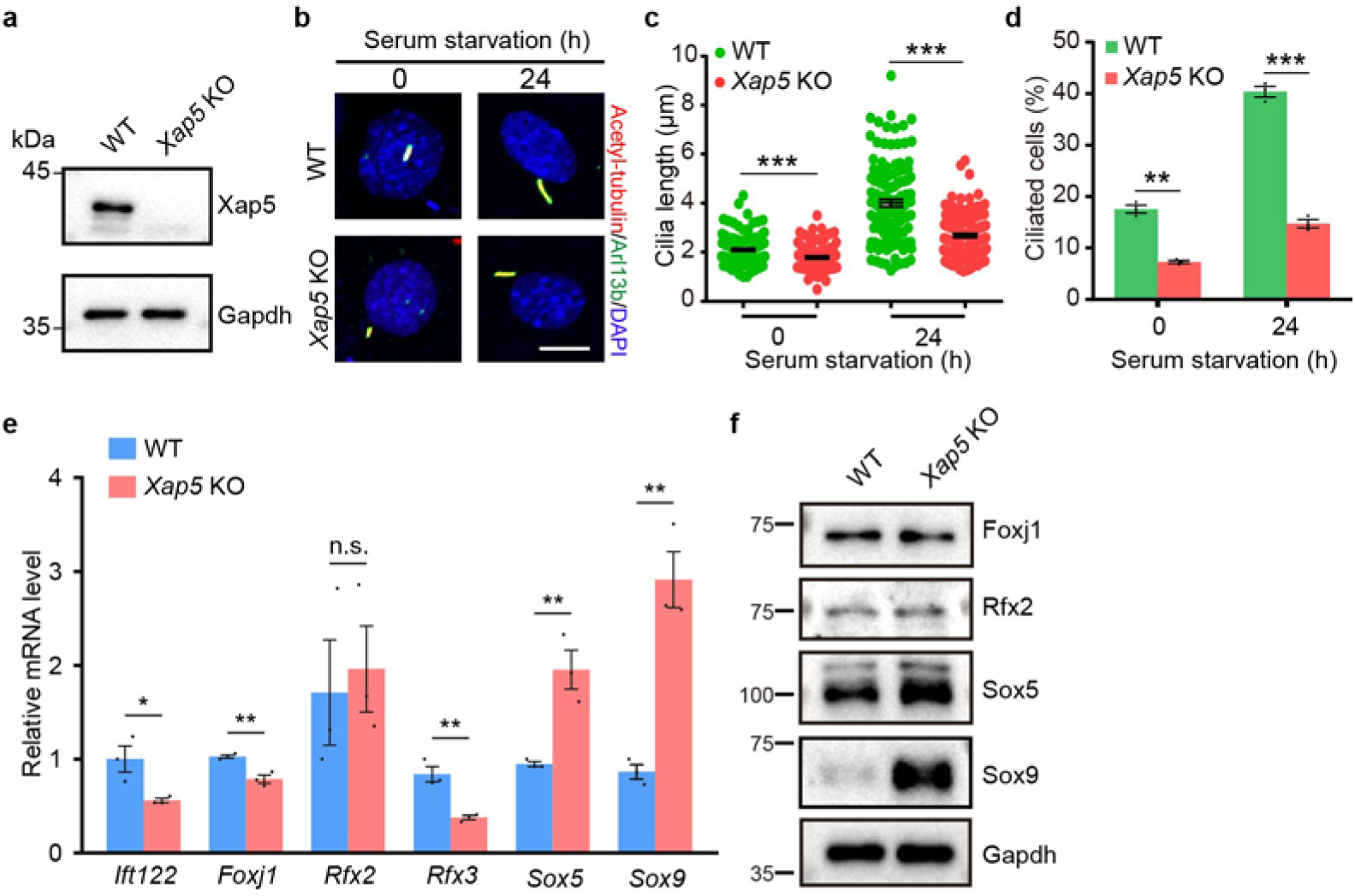
Xap5 loss reduces primary cilium length and ciliation frequency in mIMCD-3 cells. **a** Western blot analysis of Xap5 in WT and *Xap5* KO cells. **b** Representative immunofluorescence images of primary cilia before (0 h) and after 24 h of serum starvation. Cilia were stained for acetylated tubulin (red) and Arl13b (green). Nuclei were counterstained with DAPI (blue). Scale bar, 10 μm. **c** Quantification of primary cilium length at the indicated time points. Data are presented as mean ± SEM (n = 3 independent experiments). **d** Quantification of the percentage of ciliated cells as in **c**. Data are presented as mean ± SEM (n = 3 independent experiments). **e** RT– qPCR analysis of the indicated gene levels in WT and *Xap5* KO cells after 24 h of serum starvation, normalized to *Gapdh* and expressed relative to one WT sample. Data are presented as mean ± SEM from three independent experiments. **f** Representative western blots of the indicated proteins in WT and *Xap5* KO cells after 24 h of serum starvation. Gapdh serves as a loading control, and blots are representative of three independent experiments in **a** and **f**. All statistical significance between WT and KO was determined by multiple two-tailed unpaired t-tests. n.s., not significant; \**P* < 0.05; \*\**P* < 0.01; \*\*\**P* < 0.001. Source data are provided as a Source Data file.

We next asked whether the expression changes identified in NIH/3T3 cells were shared by this model. As in NIH/3T3 cells, *Rfx3* and *Ift122* mRNA levels were significantly lower in *Xap5* KO mIMCD-3 cells than in WT cells. *Foxj1* mRNA levels, which were unchanged in NIH/3T3 cells, were also lower, and *Rfx2* mRNA levels did not differ significantly between genotypes. By contrast, *Sox5* and *Sox9* mRNA levels were higher in *Xap5* KO mIMCD-3 cells (Fig. 8e). Immunoblotting detected Foxj1 and Rfx2 in both genotypes and showed higher Sox5 and Sox9 protein levels in *Xap5* KO cells (Fig. 8f). Thus, Xap5 loss shortened primary cilia and lowered Rfx3 and Ift122 expression in both cell models, whereas the Sox5 and Sox9 responses were opposite. Together with the ciliary defects, these results extend the role of Xap5 in primary ciliogenesis and ciliogenic gene expression from fibroblasts to renal epithelial cells.

## Discussion

The assembly of the cilium, a highly conserved organelle, requires a sophisticated and hierarchical transcriptional program that is responsive to extracellular cues and precisely adapted to generate enormous functional diversity^13,19-22,29^. The transcriptional program’s complexity is evident in its role in regulating flagella in unicellular organisms like *Chlamydomonas reinhardtii* and orchestrating the synchronous formation of hundreds of motile cilia in the mammalian airway^14,43-46^. In mammals, a core ciliogenic program is controlled by the Rfx family, which act as master architects for the core ciliary structure^20,23,47-49^. Built upon this foundation are multitiered regulatory cascades, often initiated by morphogenetic signaling pathways, that deploy cell-type-specific “selector cassettes”^14,20^. For the highly specialized motile cilia, this cascade involves key initiators like GMNC/MCIDAS (antagonized by GMNN)^50-53^, integrates context-specific factors like NOTO^54,55^, and converges on downstream effectors TAp73/MYB^56-58^, which in turn activate the master regulator Foxj1 to execute the motile program^27,28^. For sensory and primary cilia, genetic studies have established requirements for Rfx factors in *C. elegans*^24^ and *Drosophila*^25^, and for Rfx4 in the developing mouse nervous system^26^. Beyond these established roles, the broader upstream transcriptional hierarchy in primary-ciliated cells remains less well defined than that of the motile program.

The ancient, evolutionarily conserved transcription factor Xap5 provides an entry point for investigating this upstream regulatory hierarchy^30,59,60^. Xap5 regulates flagellar assembly in *Chlamydomonas reinhardtii*^30^ and is widely expressed in mouse tissues^31^. Our recent work revealed that during the early stage of mouse spermatogenesis, Xap5 functions upstream of Rfx2 and Foxj1 to activate ciliary gene expression^31^. These observations raised the central question addressed in this study: does this same factor also act in cells that form primary cilia and, if so, through which downstream effectors?

In this study, we show that Xap5 is required for normal primary cilia in two mouse cell models: its loss shortened primary cilia in NIH/3T3 fibroblasts and reduced both ciliation frequency and cilium length in mIMCD-3 renal epithelial cells (Figs. 1e, f and 8b-d). In NIH/3T3 cells, Xap5 associates with the nuclear protein Nono, and both proteins are required for normal cilium length and for normal expression of a shared set of cilia-associated genes that includes the transcription factor genes *Sox5* and *Sox9*, whose promoters are occupied by both Xap5 and Nono; we refer to this regulatory pair as the Xap5–Nono module (Figs. 1-5). Sox5, a target of this module, is itself required for normal cilium length (Fig. 6). Across the two cell models, Xap5 loss lowered *Rfx3* and *Ift122* mRNA levels in both, whereas Sox5 and Sox9 showed opposite responses at both the mRNA and protein levels, decreasing in NIH/3T3 cells and increasing in mIMCD-3 cells (Fig. 4d, f; Supplementary Fig. 3e; Fig. 8e, f). The ciliary phenotypes in both models, together with the regulation of multiple ciliogenic transcription factors and a broader cilia-associated gene program, identify Xap5 as a master regulator of primary cilia formation. Together with our earlier findings in spermatogenesis^31^, these results support a model in which Xap5’s conserved upstream regulatory capacity is repurposed through downstream effectors and transcriptional outputs that vary with cellular context.

Specifically, we find that the Xap5–Nono module co-regulates Sox5 and Sox9 in NIH/3T3 cells, positioning them downstream of the module. The essential role of Sox9 in promoting primary cilia formation is well-established in multiple developmental contexts^34-36^. Significantly, our data identify Sox5 as a previously unrecognized regulator of primary cilium length: Sox5 loss shortened cilia after serum starvation and lowered the expression of cilia-related genes that were also lowered by loss of Xap5 or Nono (Figs. 6 and 7). This organization is consistent with the “selector cassette” concept^20^, in which context-specific deployment of transcription factors modifies the execution of a conserved program; we therefore propose that Sox5 and Sox9 function as such a cassette downstream of the Xap5–Nono module in NIH/3T3 cells. This proposal is consistent with evidence that Sox5 can cooperate with Sox9 in other developmental programs, such as chondrogenesis^40^, and that the short isoform S-SOX5 regulates axonemal genes in motile-cilia contexts^61,62^. In mIMCD-3 cells, by contrast, Xap5 loss increased Sox5 and Sox9 mRNA and protein levels, while the ciliary defects were accompanied by lower *Rfx3*, *Foxj1* and *Ift122* mRNA levels (Fig. 8b–f). Together, these findings support a model in which the Xap5–Nono module acts upstream of Sox5 and Sox9 to organize a ciliogenic transcriptional program.

Our results suggest a framework in which Xap5 occupies sites with a similar sequence preference in different cellular contexts, whereas its transcriptional outputs depend on the context. Xap5 peaks in NIH/3T3 cells and in mouse testis^31^ shared a similar GC-rich motif (Fig. 5b). In NIH/3T3 peak windows, motif matches occurred more frequently and their highest-scoring sites were more concentrated near summits than in the specified dinucleotide-preserving shuffled references, supporting a link between GC-rich sequences and Xap5 occupancy (Supplementary Fig. 5a-c). Transcriptional responses, however, differed between contexts: the *Rfx2* promoter is occupied by Xap5 in both, whereas *Rfx2* mRNA levels decreased after Xap5 loss in testis^31^ and increased in NIH/3T3 cells (Supplementary Fig. 3e). In NIH/3T3 cells, core-promoter motif presence was not significantly associated with expression-response category (Supplementary Data 5). Within NIH/3T3 cells, loss of Nono lowered Sox5 and Sox9 expression while Xap5 remained enriched at their assayed promoter regions, supporting a role for Nono in sustaining transcriptional output at occupied loci (Figs. 4e, f and 5f). Associated proteins such as Nono may therefore provide one route by which occupancy is coupled to context-dependent outputs, and other partners may act similarly in other cell types. An informative comparison is provided by the C/T-rich C-box, which acts together with X-box elements to enhance DAF-19/Rfx-dependent ciliary gene expression in *C. elegans*^24,48,63^. The partial resemblance between the reverse-complemented C-box core and the GC-rich Xap5-associated motif, together with the broad conservation of the Xap5 family^30,59,60^, raises the testable possibility that Xap5-family proteins participate in related cis-regulatory architectures across lineages. Such an arrangement would be consistent with models of evolutionary co-option^1-3,8-10^, in which conserved regulators are deployed within different cis-regulatory and transcriptional contexts to generate distinct outputs.

Collectively, the discovery of this adaptable, Xap5-driven regulatory network offers new perspectives on human ciliopathies^15^. Xap5 loss impaired ciliogenesis in renal epithelial cells (Fig. 8b–d), and the program downregulated after loss of Xap5, Nono or Sox5 in NIH/3T3 cells includes Pkd2 (Figs. 4d, e and 7d), whose human orthologue is implicated in autosomal dominant polycystic kidney disease^64^. Loss of Sox9 is linked to ciliary defects and cystic epithelial phenotypes in liver and pancreas^34,35^. In the developing pancreas, *Sox9* deletion also reduces *Pkd2* expression^35^. The renal epithelial phenotype and the shared reduction in Pkd2 expression in NIH/3T3 cells provide a basis for testing whether altered Xap5-dependent gene regulation contributes to cystic epithelial disease. As upstream regulators of primary ciliogenesis continue to emerge—including the metazoan-specific SP5/SP8 factors^65,66^—our work highlights a distinct regulatory layer rooted in an evolutionarily ancient transcription factor^30,59,60^. Our previous evidence placing Xap5 upstream of Foxj1 and Rfx2 was obtained during mouse spermatogenesis^31^; whether the same regulatory hierarchy operates in somatic cells bearing motile cilia remains to be established. Future work should define the upstream cues that govern Xap5 activity and the mechanisms linking its context-dependent transcriptional outputs to ciliary phenotypes. Our results establish Xap5 as a master regulator of primary cilia formation and place the Xap5–Nono module upstream of Sox5 and Sox9 in NIH/3T3 cells. More broadly, these findings suggest that an evolutionarily ancient regulator can be repurposed through shared and context-specific downstream programs to generate distinct ciliogenic outputs.

## Materials and Methods

### Antibodies

The following primary antibodies were used in this study: from ABclonal Biotechnology, anti-Flag (IF 1:1,000; AE005), anti-Gapdh (IB 1:10,000; AC002), anti-Nono (IB 1:1,000, IF 1:500; A3800), and anti-Sox5 (IB 1:1,000; A23127); from Sangon Biotech, anti-Foxj1 (IB 1:1,000; D360355); from Solarbio, anti-Rfx2 (IB 1:1,000; K110716P); from Sigma-Aldrich, anti-Acetyl-α-Tubulin (IF 1:500; T7451); from Proteintech, anti-Arl13b (IF 1:500; 17711-1-AP); from HUABIO, anti-Sox9 (IB 1:1,000; ET1611-56); and from Prospertech, anti-Xap5 (IB 1:1,000, IF 1:500; HU-412003).

The following secondary antibodies were used: Alexa Fluor 488- and 594-conjugated goat anti-mouse/rabbit IgG (IF 1:500; ThermoFisher Scientific); HRP-conjugated goat anti-mouse IgG (IB 1:2,000; Bio-Rad, #1706516) and HRP-conjugated goat anti-rabbit IgG (IB 1:2,000; Bio-Rad, #1706515).

### Cell culture

The NIH/3T3 mouse embryonic fibroblast cell line (ATCC, CRL-1658) was cultured in Dulbecco’s Modified Eagle’s Medium (DMEM) supplemented with 10% calf serum (CS) and 1% penicillin-streptomycin (all from Life Technologies). The mIMCD-3 mouse renal epithelial cell line (ATCC, CRL-2123) was cultured in DMEM/F12 supplemented with 10% fetal bovine serum (FBS) and 1% penicillin-streptomycin (all from Life Technologies). Cells were maintained in a humidified incubator at 37°C and 5% CO□.

### Plasmid construction, transfection and generation of knockout cell lines

For protein expression, the coding sequences of mouse Xap5 and Nono were amplified from NIH/3T3 cDNA and cloned into the pECMV-MCS-Flag and pCMV-C-GFP vectors, respectively, to generate constructs encoding C-terminally tagged proteins. Site-directed mutagenesis to disrupt the Xap5 nuclear localization signal (NLS) was performed using the Mut Express II Fast Mutagenesis Kit V2 (Vazyme, C214). For CRISPR/Cas9-mediated gene knockout in NIH/3T3 cells, the all-in-one plasmid pU6-sgRosa26-1_CBh-Cas9-T2A-BFP (a gift from Ralf Kuehn; Addgene plasmid #64216) was used as the backbone. Gene-specific sgRNA sequences were inserted by replacing the original sgRosa26-1 guide RNA sequence via site-directed mutagenesis. The integrity of all constructs was confirmed by Sanger sequencing. All primer and sgRNA sequences are listed in Supplementary Tables 1 and 2, respectively.

For transient expression experiments, cells were transfected with the indicated DNA constructs using Lipofectamine 3000 (Life Technologies) according to the manufacturer’s protocol. For the rescue experiments, *Xap5* KO NIH/3T3 cells were transfected with constructs encoding GFP–Flag, WT Xap5-Flag or the NLS-mutant Xap5-Flag (ΔNLS–Flag) protein. GFP- Flag was used as an expression control in both WT and *Xap5* KO cells to assess the effects of expressing an unrelated Flag-tagged protein on ciliary length and ciliation frequency.

Stable knockout cell lines were generated by electroporation. Briefly, 1×10^6^ cells were resuspended in 250 µL of Opti-MEM I Reduced Serum Medium (Gibco). For NIH/3T3 cells, the suspension contained 20 μg of the appropriate all-in-one CRISPR–Cas9 plasmid. For mIMCD-3 cells, the suspension contained 4 μg Cas9 protein (GenScript, Z03702) and 50 pmol synthetic Xap5-targeting sgRNA (sequences are listed in Supplementary Table 2). The cell suspension was transferred to a 0.4 cm gap electroporation cuvette (Bio-Rad) and electroporated using a Gene Pulser Xcell Electroporation System (Bio-Rad) with a single exponential decay pulse (250 V, 950 µF, ∞ Ω resistance). Twenty-four hours after electroporation, individual cells were manually isolated and seeded into separate wells of a 96-well plate to establish monoclonal colonies. For NIH/3T3 cells, cells exhibiting bright BFP fluorescence were identified by fluorescence microscopy for clonal isolation. The resulting colonies were subsequently expanded and validated for successful gene knockout.

### Immunofluorescence and microscopy

To induce primary cilia formation, NIH/3T3 and mIMCD-3 cells were seeded onto sterile, poly-D-lysine-coated (Gibco) glass coverslips (CITOTEST). After 24 h in growth medium, cells were switched to serum-free medium for 24 or 48 h (NIH/3T3) or 24 h (mIMCD-3). The 0 h samples were collected before serum withdrawal. For rescue experiments, NIH/3T3 cells were transfected 18 h after seeding. The medium was replaced with serum-free medium 6 h after transfection, and cells were serum-starved for 24 h.

Cells on coverslips were washed three times with PBS and fixed with 4% paraformaldehyde (PFA) in PBS for 10 min at room temperature. Following fixation, residual PFA was quenched with 50 mM NH_4_Cl in PBS for 5 min. Cells were then permeabilized with 0.2% Triton X-100 for 15 min and blocked with 3% bovine serum albumin (BSA) in PBS for 1 h. Primary antibody incubations were performed for 1 h at room temperature in a humidified chamber. After washing, cells were incubated with Alexa Fluor-conjugated secondary antibodies for 1 h, followed by a 5 min counterstain with DAPI (Sigma-Aldrich). Finally, coverslips were washed and mounted onto glass slides using Fluoromount-G mounting medium (SouthernBiotech). This protocol was used for all cilia detection and Xap5 ectopic expression experiments. For the detection of ectopically expressed Nono-GFP, cells were fixed and washed as above, then directly counterstained with DAPI before mounting, omitting the permeabilization and antibody incubation steps.

General fluorescence for confirming ectopic protein expression was assessed using a Zeiss Axio Vert. A1 inverted microscope. High-resolution imaging of primary cilia was performed on a Leica SP8 confocal microscope using a 63× oil-immersion objective. The percentage of ciliated cells and cilia length were quantified from confocal images using the Automated Cilia Detection and Counter (ACDC) software (v0.93)^67^. The nuclear area was quantified from DAPI-stained images using Fiji software.

### RNA isolation and RT-qPCR

Total RNA was isolated from cells using the FastPure Cell/Tissue Total RNA Isolation Kit (Vazyme, RC112) according to the manufacturer’s protocol. For each sample, 500 ng of total RNA was reverse-transcribed into cDNA using the HiScript II 1st Strand cDNA Synthesis Kit (Vazyme, R212). RT-qPCR was then performed with ChamQ SYBR qPCR Master Mix (Vazyme, Q311) on an Applied Biosystems StepOnePlus Real-Time PCR System. Relative gene expression was calculated using the 2^−ΔΔCt^ method, with *Gapdh* serving as the internal normalization control. All primer sequences are listed in Supplementary Table 1.

### Western blotting

Cells were lysed on ice in RIPA buffer (Beyotime, P0013B) supplemented with 1× protease inhibitor cocktail (Roche, 11873580001) and 1 mM PMSF (Sigma, 93482). The resulting lysates were resolved by SDS-PAGE and transferred to polyvinylidene fluoride membranes (Millipore, IPVH00010) using a wet transfer system (Tanon). Membranes were blocked with 5% non-fat milk in TBST (20 mM Tris-HCl, pH 7.5, 150 mM NaCl, 0.1% Tween-20) for 1 h at room temperature, followed by incubation with primary antibodies overnight at 4 °C. After washing, membranes were incubated with HRP-conjugated secondary antibodies for 1 h at room temperature. Chemiluminescence signals were developed using the ECL Super Kit (ABClonal, RM02867) and visualized with a Tanon 5200 Chemiluminescence Imaging System.

### Co-immunoprecipitation (Co-IP) and mass spectrometry

For Co-IP and IP-MS experiments, cultured cells were washed twice with cold 1× PBS and harvested by scraping. Total proteins were extracted on ice using a Co-IP buffer containing 50 mM HEPES (pH 7.5), 200 mM KCl, 1 mM EGTA, 1 mM MgCl_2_, 10% (v/v) glycerol, 0.5 mM DTT, 0.3% (v/v) NP-40, and 1× protease inhibitor cocktail (Roche). For immunoprecipitation, antibodies were first crosslinked to Protein A/G magnetic beads (Invitrogen, 80105G) using 5 mM BS^3^. The resulting antibody-conjugated beads were then incubated with the cell lysates for 2 h at 4°C with gentle rotation. After extensive washing, bound proteins were eluted with 0.1 M Glycine-HCl (pH 2.5). For western blot analysis, the eluates and input samples were neutralized, mixed with SDS loading buffer, boiled for 5 min, and subsequently analyzed by immunoblotting. For protein identification, the eluates were sent for analysis by liquid chromatography with tandem mass spectrometry (LC-MS/MS) at Novogene (Beijing, China). The complete list of identified proteins is provided in Supplementary Data 1.

### RNA-sequencing and analysis

For transcriptomic analysis, cells were plated for 24 h and subsequently serum-starved for 24 h. At the end of the starvation period, cells were immediately washed three times with 1× PBS and then lysed directly in TRIzol Reagent (Invitrogen) for total RNA extraction. Library construction and paired-end (2 × 150 bp) sequencing on an Illumina NovaSeq 6000 platform were performed by BioMarker Technologies (Beijing, China).

The raw sequencing reads were mapped to the Mus musculus reference genome (GRCm39/mm39) using HISAT2^68^ (v2.1.0). StringTie^69^ (v1.3.3b) was subsequently used to assemble the mapped reads. Gene expression levels were quantified as fragments per kilobase of transcript per million fragments mapped (FPKM). Differential expression analysis between experimental groups was performed using DESeq2^70^ (v1.12.4). Genes with a false discovery rate (FDR) < 0.01 and an absolute fold change ≥ 1.5 were considered significantly differentially expressed. Gene Ontology (GO) enrichment analysis was performed using the DAVID database. Gene Set Enrichment Analysis (GSEA) was conducted using the OECloud platform (https://cloud.oebiotech.com).

### CUT&Tag-sequencing and analysis

For CUT&Tag analysis, NIH/3T3 cells were plated for 24 h and subsequently serum-starved for 24 h. Following harvesting, 100,000 cells were collected for each sample. Library construction was performed using the Hyperactive Universal CUT&Tag Assay Kit for Illumina Pro (Vazyme, TD904) according to the manufacturer’s protocol. The resulting libraries were sequenced on an Illumina NovaSeq platform (150 bp paired-end reads) at Novogene Co., Ltd. (Beijing, China).

For bioinformatic processing, raw reads were assessed for quality with FastQC^71^ (v0.11.9) and trimmed with fastp^72^ (v0.20.0; parameters: --length_required 15, --n_base_limit 6). Clean reads were aligned to the Mus musculus reference genome (mm39) using BWA-MEM (v0.7.12; parameters: -k 32, -T 30, -M). Peak calling was performed using MACS2^73^ (v2.1.0) with a q-value threshold of 0.05 and the following parameters: -f AUTO --call-summits --nomodel –shift -100 --extsize 200 --keep-dup all. For *de novo* motif discovery, peak summits were extended to 500 bp and analyzed using the HOMER^74^ (v4.9.1) findMotifsGenome.pl script. Peak annotation and association with the nearest genes were performed using the ChIPseeker^75^ R package. Genome browser tracks were generated using the Integrative Genomics Viewer (IGV, v2.18.1)^76^.

Row-normalized Xap5 motifs from NIH/3T3 cells and mouse testis were compared using TOMTOM v5.5.9 (Pearson distance; reverse-complement matching) against 879 JASPAR2024 CORE vertebrate matrices supplemented with the other motif. Similarity required reciprocal P < 0.05 and alignment across all ten positions of the shorter motif. Known-factor searches used JASPAR2024 and HOCOMOCO v14, retaining q < 0.05 matches and the top ten by P value. Core promoters were defined on GRCm39/GENCODE M36 as −500 to +250 bp (751 bp) around representative TSSs, prioritizing canonical transcripts, then greatest total non-overlapping exonic length, with transcript ID resolving ties. The original NIH/3T3 matrix was scanned on both strands using HOMER v5.1 at 8.034194 nats. In the eleven-gene composition-control analysis, maximum continuous PWM scores (bits) were compared with 1,000 dinucleotide-preserving shuffles per promoter, holding non-ACGT positions fixed. Empirical P values were calculated as (1 + number of shuffled maxima ≥ the observed maximum)/1,001 and Benjamini–Hochberg-adjusted across the eleven genes.

Summit analyses used 1,001-bp windows around 45,495 eligible WT NIH/3T3 Xap5 peaks and 200 whole-window and 200 local-bin dinucleotide-preserving shuffles per window. A custom scanner applied the original matrix and threshold to both strands, retaining highest-scoring passing sites with equal weights for exact ties. Q was the fraction of hit-positive windows; L, central (±50 bp) highest-match weight divided by the number of hit-positive windows; and p, the same central weight divided by all windows. Control ratios used pooled counts and weights across window–shuffle pairs; comparisons were descriptive. Peak labels required ≥1-bp core overlap; joint labels required motif and peak within the same core, irrespective of mutual overlap. Sox5/Sox9 amplicons were mapped by exact primer matching to both strands.

Expression associations used a separate RNA-seq reanalysis of WT versus Xap5 KO NIH/3T3 cells after 24 h of serum starvation. UP and DOWN genes had adjusted P < 0.01 and log2 fold changes ≥log2(1.5) or ≤−log2(1.5), respectively; STABLE genes had adjusted P ≥0.05 and absolute log2 fold changes <log2(1.2). Logistic regression in R v4.3.3 modelled core-promoter motif presence by expression category (STABLE reference) in 7,068 uniquely mapped genes with complete covariates. GC fraction, CpG density and log(1 + WT mean normalized count) each used three-degree-of-freedom natural splines. Two-sided Wald P values were Holm-adjusted across the two comparisons with STABLE, and Wald 95% confidence intervals were reported.

### CUT&Tag-qPCR

For targeted validation, DNA from 100,000 cells was prepared in parallel using the same CUT&Tag kit according to the manufacturer’s protocol. qPCR was performed using the resulting DNA. Samples processed with IgG served as negative controls, and an exogenous DNA spike-in was used as an internal normalization control. Enrichment at specific loci was calculated relative to the spike-in DNA control, and the 2^−ΔΔCt^ method was used to compare experimental samples. For comparisons of Xap5 enrichment between WT and Nono KO NIH/3T3 cells, spike-in-normalized signals were expressed as S = Ct(spike-in) − Ct(target). Log2 enrichment over IgG was calculated as S(anti-Xap5) − S(IgG), and fold enrichment as 2 raised to this value. WT and KO samples from the same independent experiment were paired for statistical analysis. All primer sequences used for CUT&Tag-qPCR are listed in Supplementary Table 1.

### Statistical analysis

Statistical analysis was performed using GraphPad Prism v6.01 or v8.0.2. All data were presented as mean ± SEM unless otherwise indicated. Values of P < 0.05 were considered statistically significant. Comparisons between independent groups were calculated using an unpaired, parametric, two-sided Student’s t-test. For targeted RT-qPCR comparisons, tests were performed on linear relative-expression values. Xap5 CUT&Tag-qPCR comparisons between WT and Nono KO NIH/3T3 cells used paired, two-sided Student’s t-tests on IgG-corrected log2 enrichment values from three matched experiments.

## Data availability

The RNA-sequencing data generated in this study have been deposited in the Gene Expression Omnibus (GEO) database under accession code GSE305355. The CUT&Tag-sequencing data reported in this paper are available in the GEO database under accession code GSE304980. All other data supporting the findings of this study are available within the article and its Supplementary Information files. Source data are provided with this paper.

## Acknowledgements

This work was supported by the National Natural Science Foundation of China (grant number: 32170702), the Natural Science Foundation for Distinguished Young Scholars of Hubei Province (grant number: 2025AFA060), the Wuhan Municipal Education Bureau’s Program for the Integration of Research and Education (grant number: 2025KCJ03), and the Jianghan University Research Fund (grant number: ZGBR202603). We thank Dr. Peter Swoboda (Karolinska Institute) for his constructive comments on the preprint version of this manuscript, specifically for pointing out the sequence similarity between the Xap5 motif and the *C. elegans* C-box.

## Author Contributions

W.W., X.Z., Y.Q., X.M., G.T. and Z.H. conceived and designed the research. W.W., X.Z., Y.Q., X.M., S.C., Y.C., S.L., W.C., J.Y., X.Y., H.L., J.X., C.X. and H.J. performed the research. W.W., G.T. and Z.H. contributed new reagents/analytic tools. W.W., H.W., G.T. and Z.H. analyzed the data. W.W. and Z.H. wrote the manuscript. All authors reviewed the manuscript and approved the final version.

## Supplementary information

**Supplementary Data 1**

Proteins identified by Xap5 IP-MS in NIH/3T3 cells

**Supplementary Data 2**

Differentially expressed genes between WT and *Xap5* KO NIH/3T3 cells identified by RNA-seq

**Supplementary Data 3**

Differentially expressed genes between WT and *Nono* KO NIH/3T3 cells identified by RNA-seq

**Supplementary Data 4**

Curated list of cilia-associated genes used in the study

**Supplementary Data 5**

Xap5 CUT&Tag peak-associated genes and motif analyses

**Supplementary Data 6**

Genes associated with Nono CUT&Tag peaks in NIH/3T3 cells

**Supplementary Data 7**

Differentially expressed genes between WT and *Sox5* KO NIH/3T3 cells identified by RNA-seq

## Supplementary Figures

**Supplementary Fig. 1.**
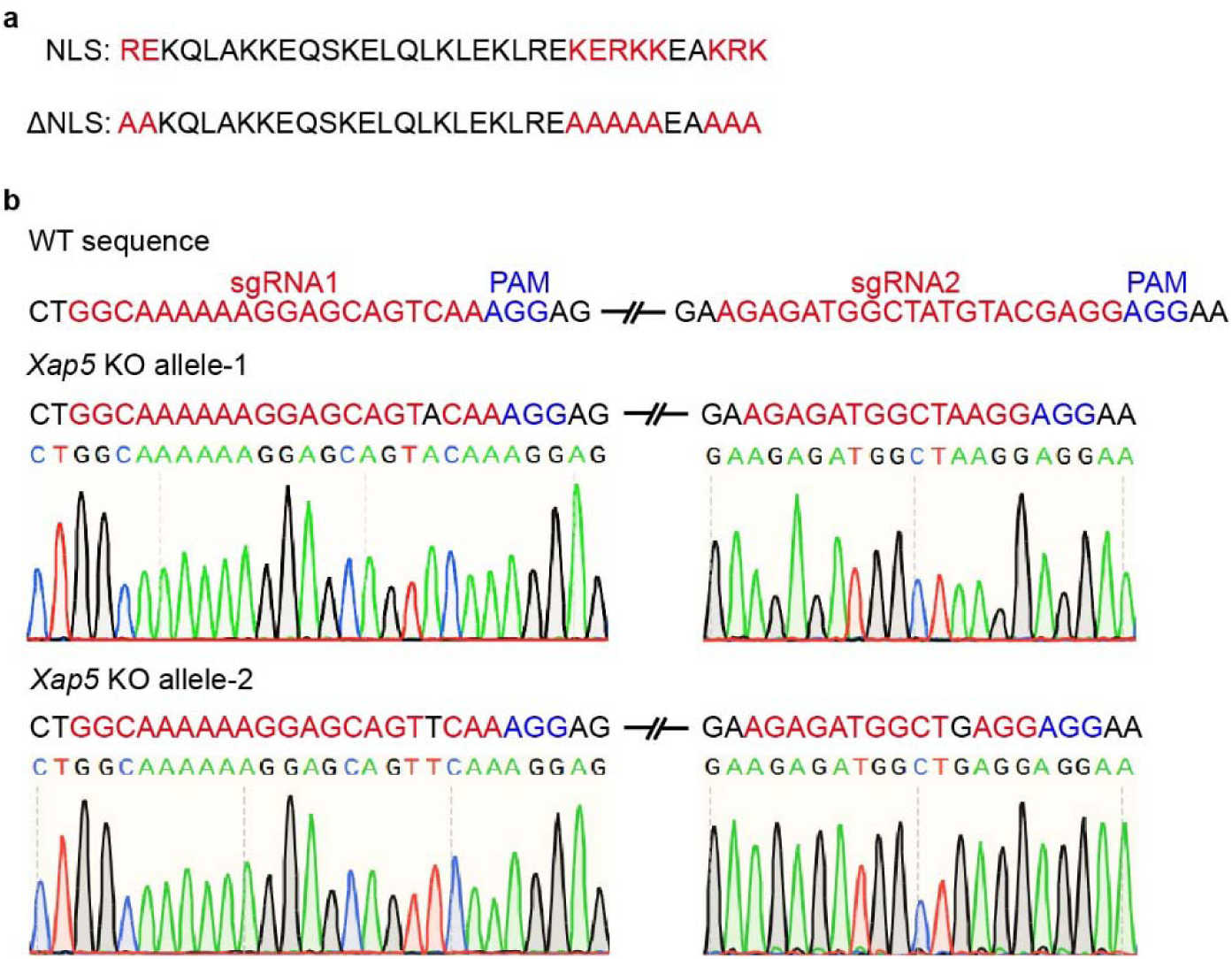
Design and validation of NLS-mutant and CRISPR/Cas9-mediated *Xap5* KO alleles. **a** Amino acid sequences of the predicted NLS region in WT Xap5 and the mutant designated ΔNLS. The substituted residues and their WT counterparts are highlighted in red. **b** Validation of CRISPR/Cas9-mediated knockout of *Xap5*. Top: Schematic of the wild-type genomic locus showing the target sites for the two sgRNAs (Xap5-sgRNA1 and Xap5-sgRNA2, red) and their adjacent PAM sequences (blue). Bottom: Sanger sequencing chromatograms from a selected knockout clone revealing two distinct mutant alleles (allele-1 and allele-2). Both sequenced alleles carry a 1-bp insertion at the sgRNA1 site, and a 6-bp in-frame deletion at the sgRNA2 site.

**Supplementary Fig. 2.**
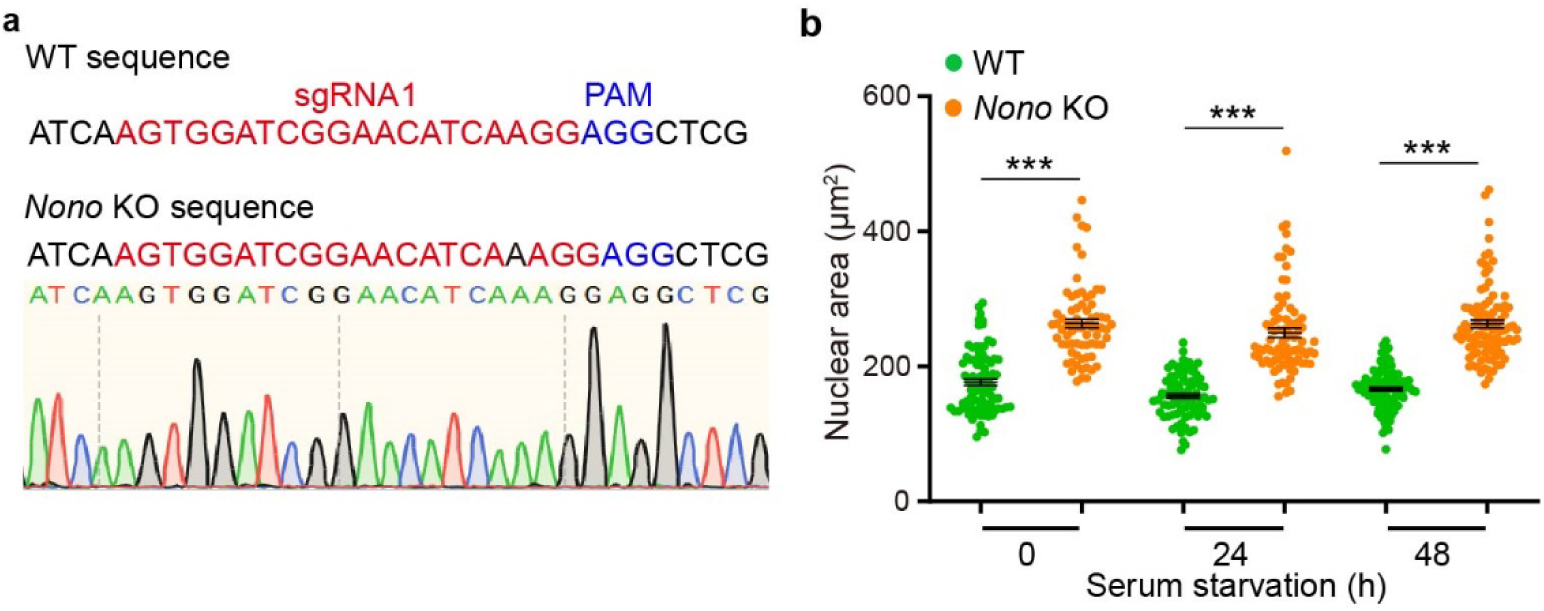
Validation of *Nono* KO and its impact on nuclear architecture. **a** Sanger sequencing of the targeted *Nono* region in the analyzed NIH/3T3 KO clone. The Nono-sgRNA1 target sequence and adjacent PAM are highlighted in red and blue, respectively. The resulting KO sequence harbors a frameshift-inducing indel at the target site, consistent with successful gene disruption. **b** Quantification of nuclear area (μm^2^) in WT and *Nono* KO cells over a 48-h time course of serum starvation. Data are presented as the mean ± SEM (n = 3 independent biological replicates). Statistical significance between WT and KO cells at each time point was determined by multiple two-tailed unpaired t-tests. \*\*\**P* < 0.001. Source data are provided as a Source Data file.

**Supplementary Fig. 3.**
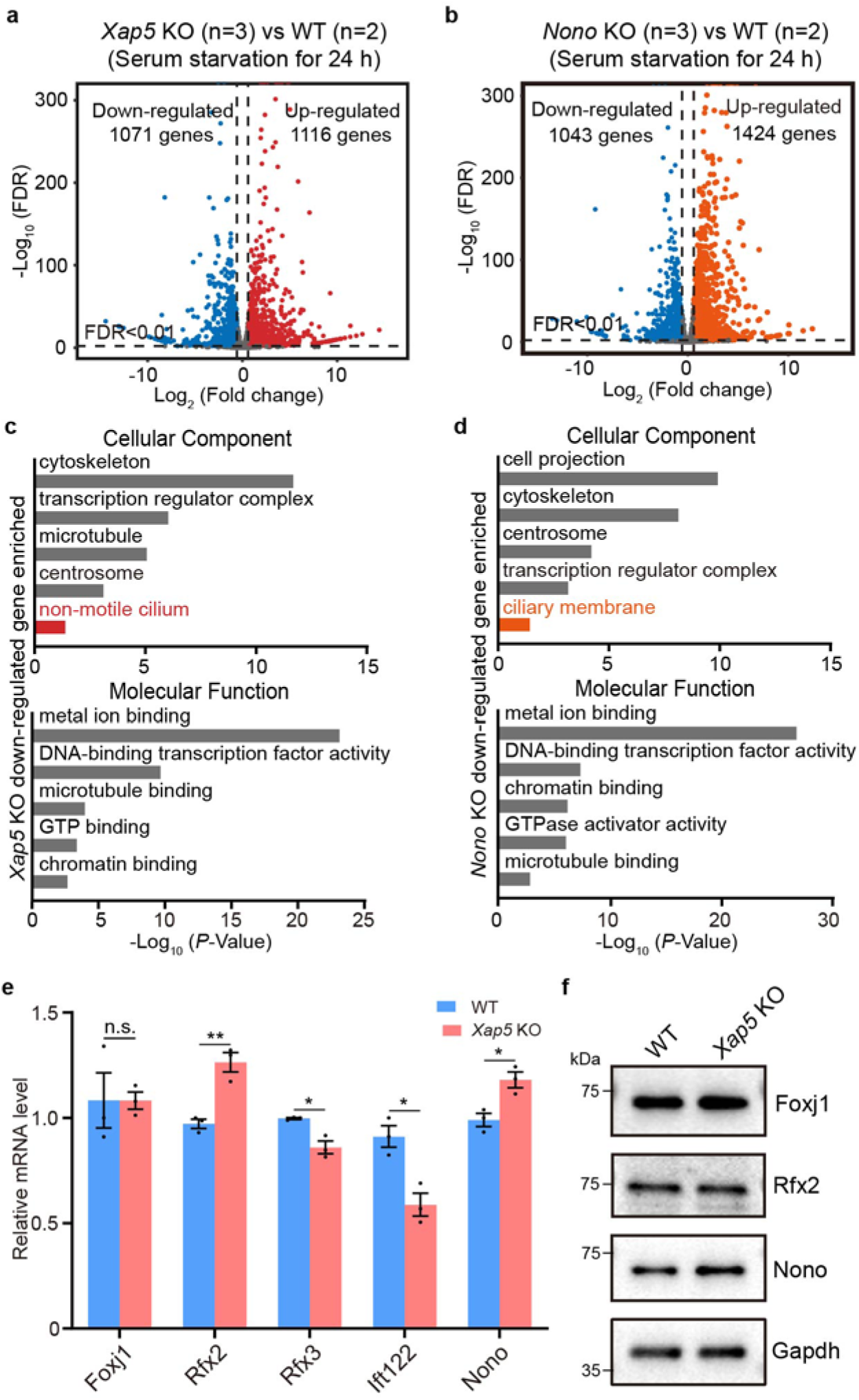
Transcriptomic profiling of *Xap5* and *Nono* KO cells. All analyses were performed on NIH/3T3 cells following 24 h of serum starvation. **a** Volcano plot of differentially expressed genes in *Xap5* KO (n=3 biological replicates) versus WT (n=2) cells. Using cutoffs of absolute fold change ≥ 1.5 and a false discovery rate (FDR) < 0.01 (highlighted dots), we identified 1,116 upregulated and 1,071 downregulated genes. **b** Volcano plot for *Nono* KO (n=3) versus WT (n=2) cells, analyzed using the same conditions and cutoffs as in **a**. This analysis identified 1,424 upregulated and 1,043 downregulated genes. **c**, **d** Representative enriched Gene Ontology (GO) terms for Cellular Component and Molecular Function categories among genes downregulated in *Xap5* KO (**c**) and *Nono* KO (**d**) cells. Cilia- and transcription-related terms are prominent in both datasets, reinforcing their dual roles in ciliogenesis and nuclear function. **e** RT-qPCR analysis of the indicated genes in WT and *Xap5* KO cells. Data are presented as mean ± SEM (n = 3 independent experiments). All statistical significance was determined by multiple two-tailed unpaired t-tests. \**P* < 0.05; \*\**P* < 0.01; n.s., not significant. **f** Western blots of Foxj1, Rfx2 and Nono in WT and *Xap5* KO cells, representative of at least three independent experiments. Gapdh served as the loading control. Source data are provided as a Source Data file.

**Supplementary Fig. 4.**
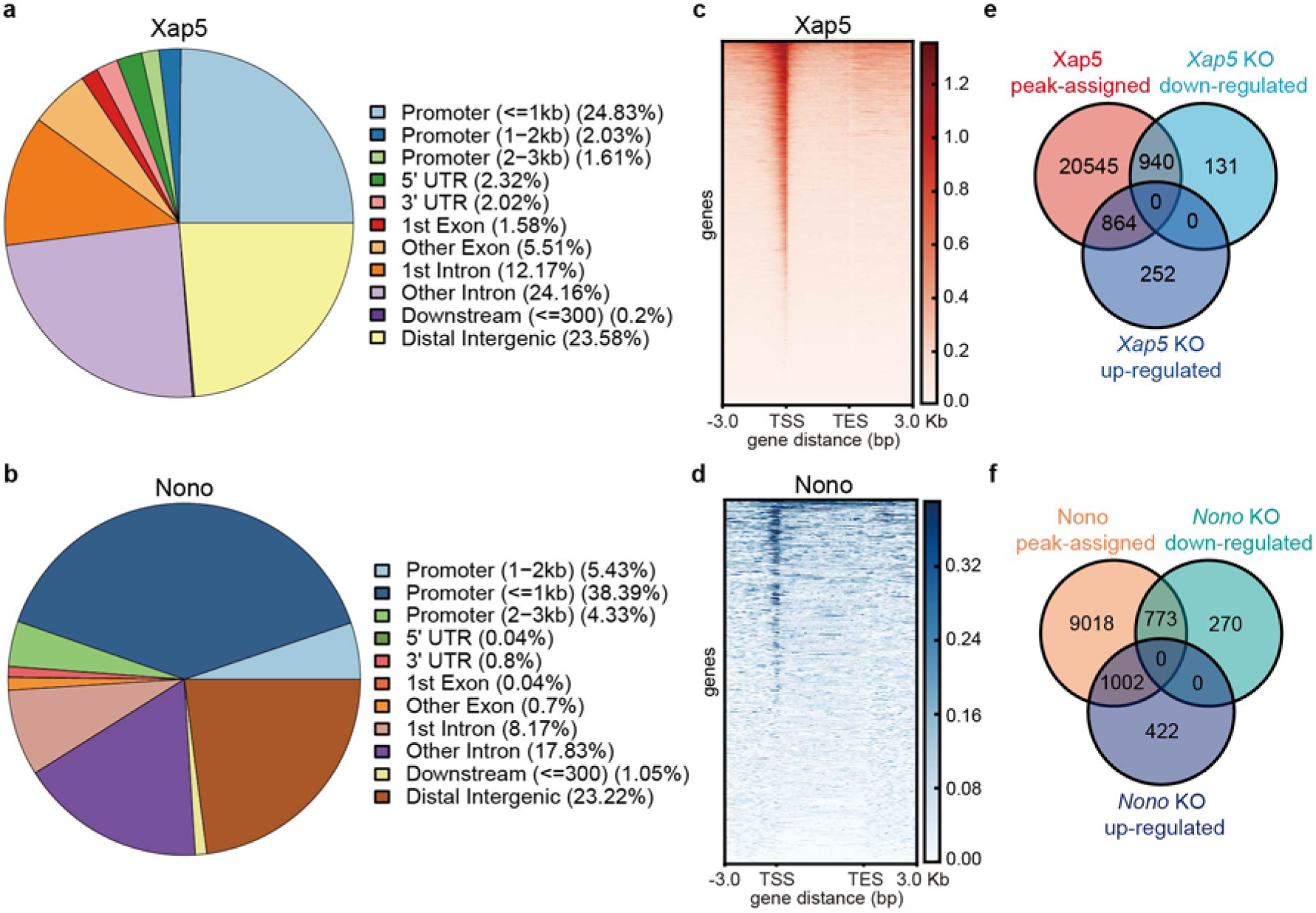
Genomic distribution of Xap5 and Nono CUT&Tag peaks and integration with RNA-seq. All analyses were performed on NIH/3T3 cells following 24 h of serum starvation. **a**, **b** Genomic distribution of CUT&Tag peaks for Xap5 (**a**) and Nono (**b**). **c**, **d** Heatmaps of Xap5 (**c**) and Nono (**d**) CUT&Tag signals across gene bodies (TSS–TES) and 3-kb flanking regions. TSS, transcription start site; TES, transcription end site. **e**, **f** Venn diagrams showing overlap between genes assigned to Xap5 (**e**) or Nono (**f**) CUT&Tag peaks and genes downregulated or upregulated in the corresponding KO cells.

**Supplementary Fig. 5.**
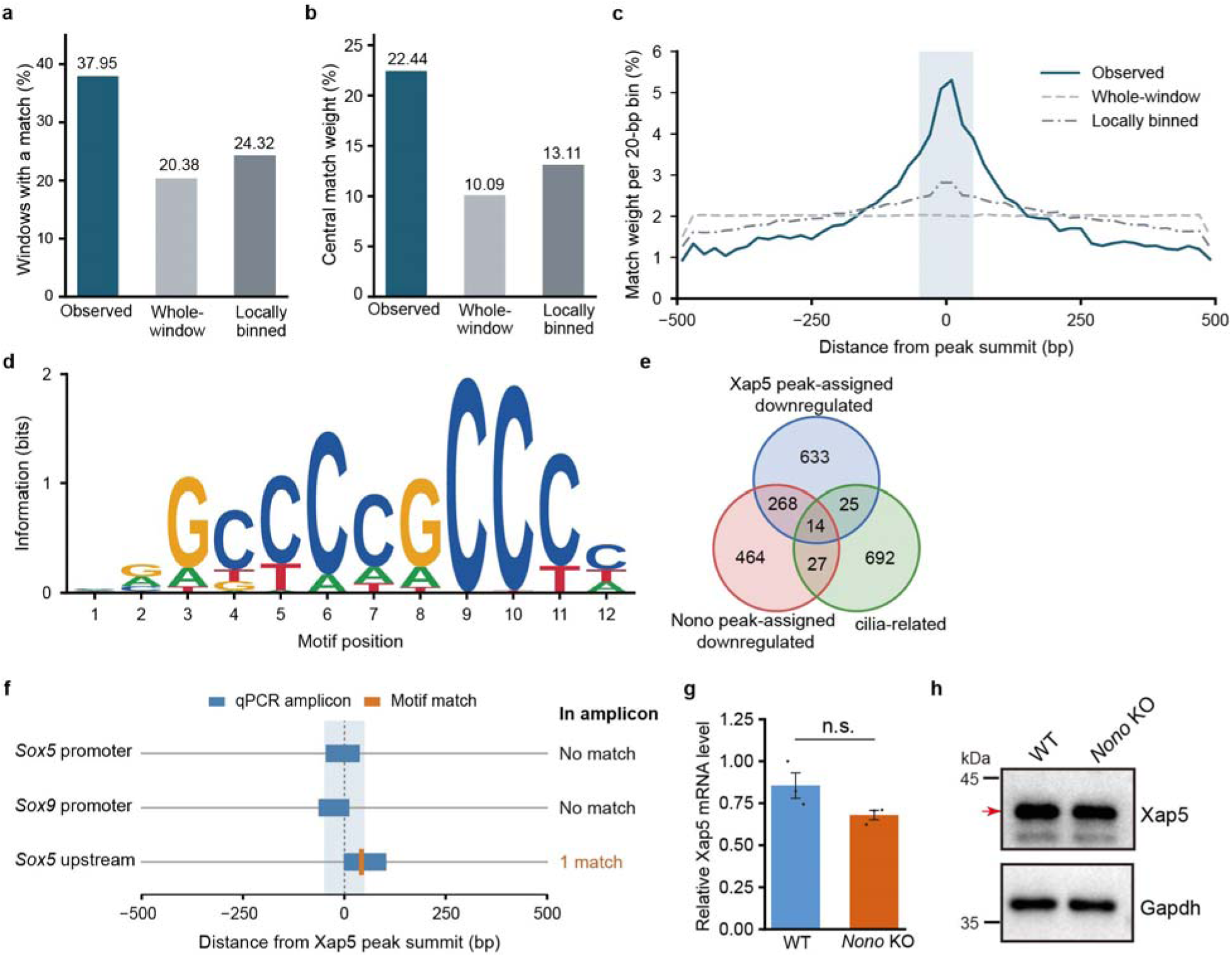
Xap5-associated motif analyses and Xap5 expression in *Nono* KO cells. **a–c** Descriptive motif analyses of 45,495 Xap5 peak windows (±500 bp), compared with pooled whole-window and locally binned dinucleotide-preserving shuffle controls (200 shuffles per window for each control). **a** Percentage of windows containing at least one motif match. **b**, **c** Percentage of highest-scoring match weight within ±50 bp of the summit (**b**) or each 20-bp bin (**c**), normalized to the total weight in match-positive windows. Each match-positive window contributes unit weight, divided equally among tied highest-scoring matches. **d** Information-content logo of the 12-position Xap5-associated motif used for scanning. **e** Overlap among Xap5 peak-assigned genes downregulated in *Xap5* KO cells, Nono peak-assigned genes downregulated in *Nono* KO cells, and the curated cilia-associated gene set (Supplementary Data 4). **f** Positions of CUT&Tag–qPCR amplicons (blue) and motif matches (orange) relative to the corresponding Xap5 peak summits. Counts refer to complete threshold-passing matches within each amplicon. Shading in **c** and **f** indicates ±50 bp. **g** RT–qPCR analysis of *Xap5* in WT and *Nono* KO NIH/3T3 cells, normalized to *Gapdh* with one WT sample as calibrator. Data are presented as mean ± SEM (n = 3 independent experiments). **h** Western blot analysis of Xap5 in WT and *Nono* KO cells. The arrow indicates Xap5; Gapdh served as a loading control. The P value in **g** was determined by a two-tailed unpaired Student’s t-test. n.s., not significant. Source data are provided as a Source Data file.

**Supplementary Fig. 6.**
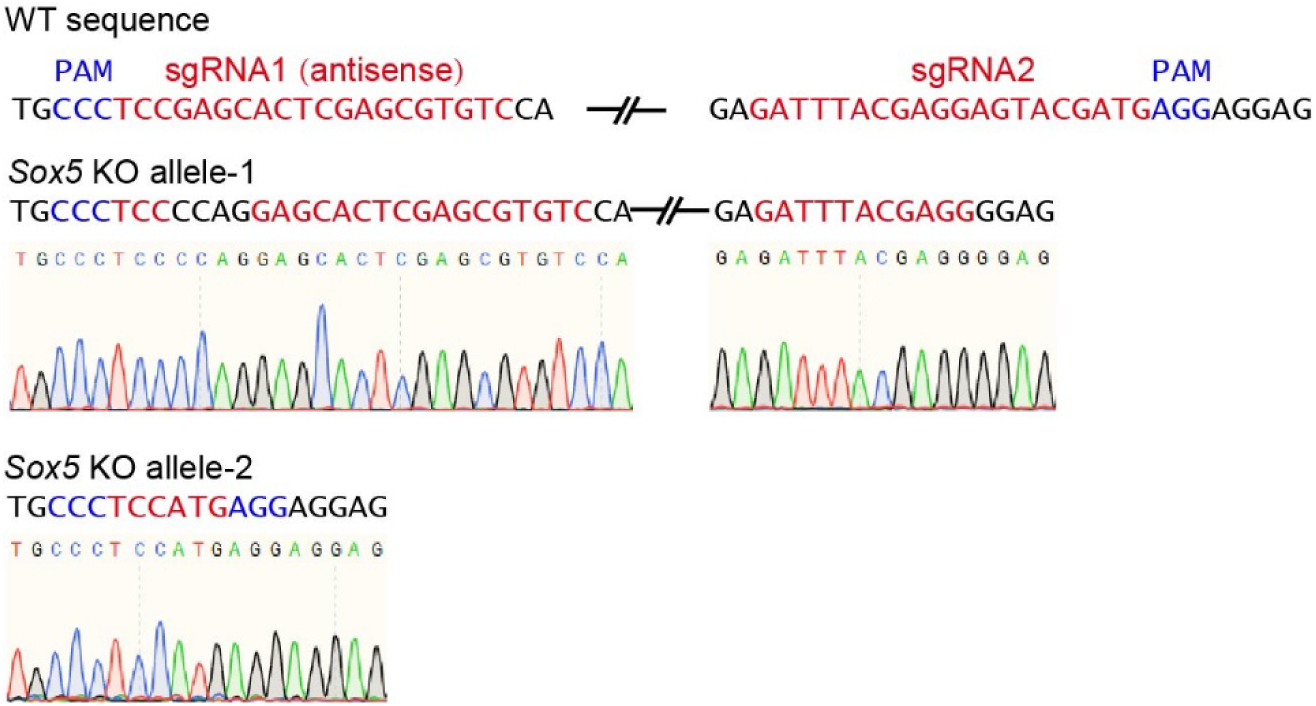
Validation of *Sox5* KO alleles by Sanger sequencing. WT sequences and Sanger sequencing chromatograms of two mutant alleles from the same NIH/3T3 KO clone are shown. The Sox5-sgRNA1 (antisense) and Sox5-sgRNA2 target sequences are highlighted in red, and PAM sequences in blue. Two independent *Sox5* KO alleles (KO allele-1 and KO allele-2) from the same clone display distinct mutations between the sgRNA cut sites. KO allele-1 harbors distinct frameshift-inducing mutations at each of the two sgRNA target sites, while KO allele-2 contains a large deletion spanning the entire region between the sgRNA sites. These results confirm successful biallelic disruption of the *Sox5* locus

**Supplementary Fig. 7.**
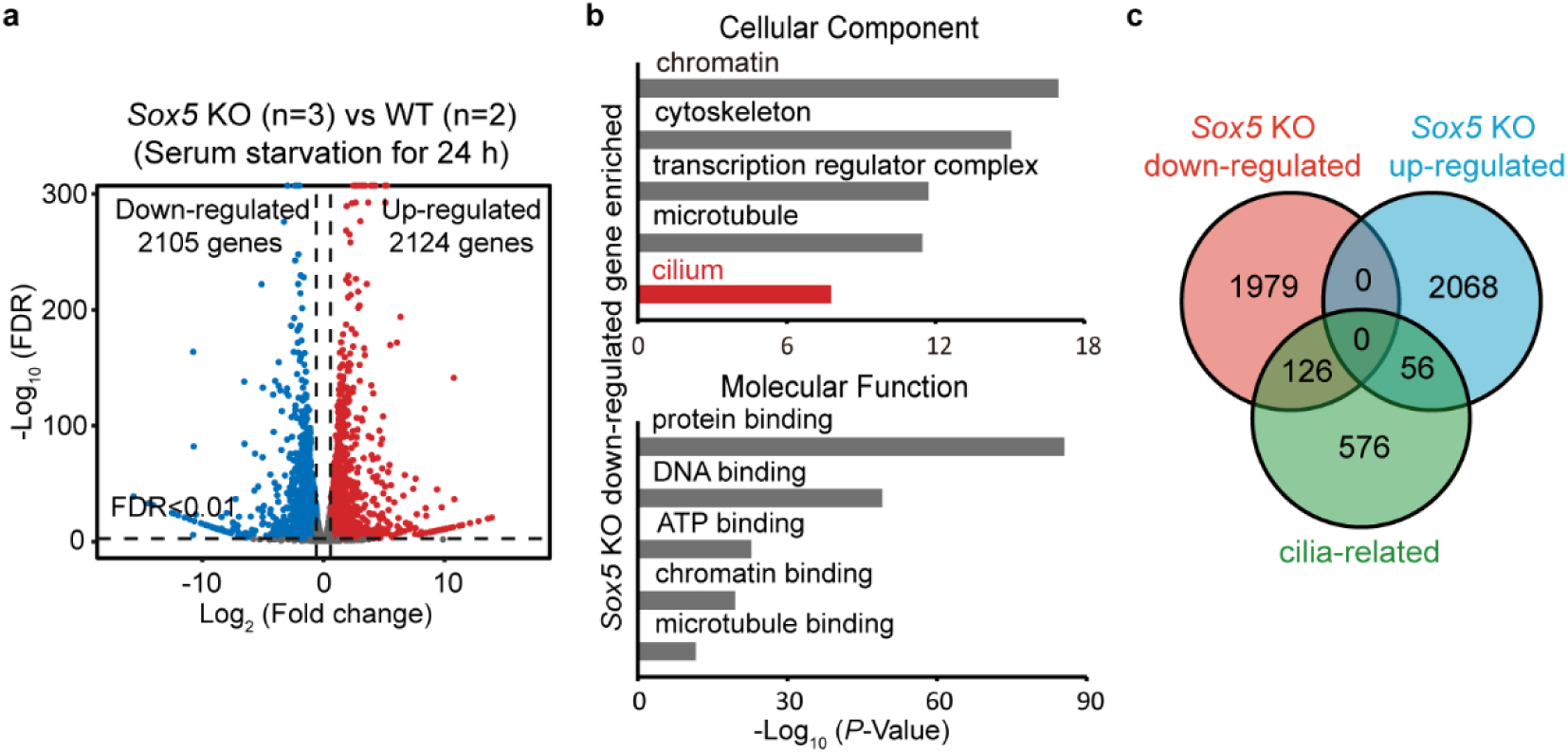
Transcriptomic analysis of *Sox5* KO cells. All analyses were performed on NIH/3T3 cells following 24 h of serum starvation. **a** Volcano plot of differentially expressed genes in *Sox5* KO (n=3 biological replicates) versus WT (n=2) cells. Using cutoffs of absolute fold change ≥ 1.5 and FDR < 0.01 (highlighted dots), we identified 2,124 upregulated and 2,105 downregulated genes. **b** Representative enriched GO terms for Cellular Component and Molecular Function categories among genes downregulated in *Sox5* KO cells. The significant enrichment of cilia-related terms reinforces the link between Sox5 and ciliogenesis. **c** Venn diagram illustrating the overlap between the curated ciliary gene set (Supplementary Data 4) and genes either upregulated or downregulated in *Sox5* KO cells.

**Supplementary Fig. 8.**
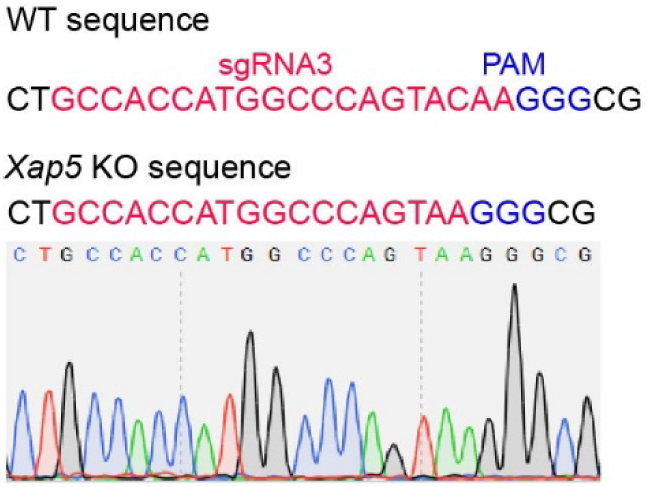
Sequence validation of *Xap5* KO in mIMCD-3 cells. Sequence analysis of the targeted *Xap5* region in WT and the analyzed *Xap5* KO mIMCD-3 clone. The Xap5-sgRNA3 target sequence and PAM sequence are highlighted in red and blue, respectively. The displayed KO sequence contains a 2-bp deletion, with the corresponding Sanger sequencing chromatogram shown below.

**Supplementary Table 1.**
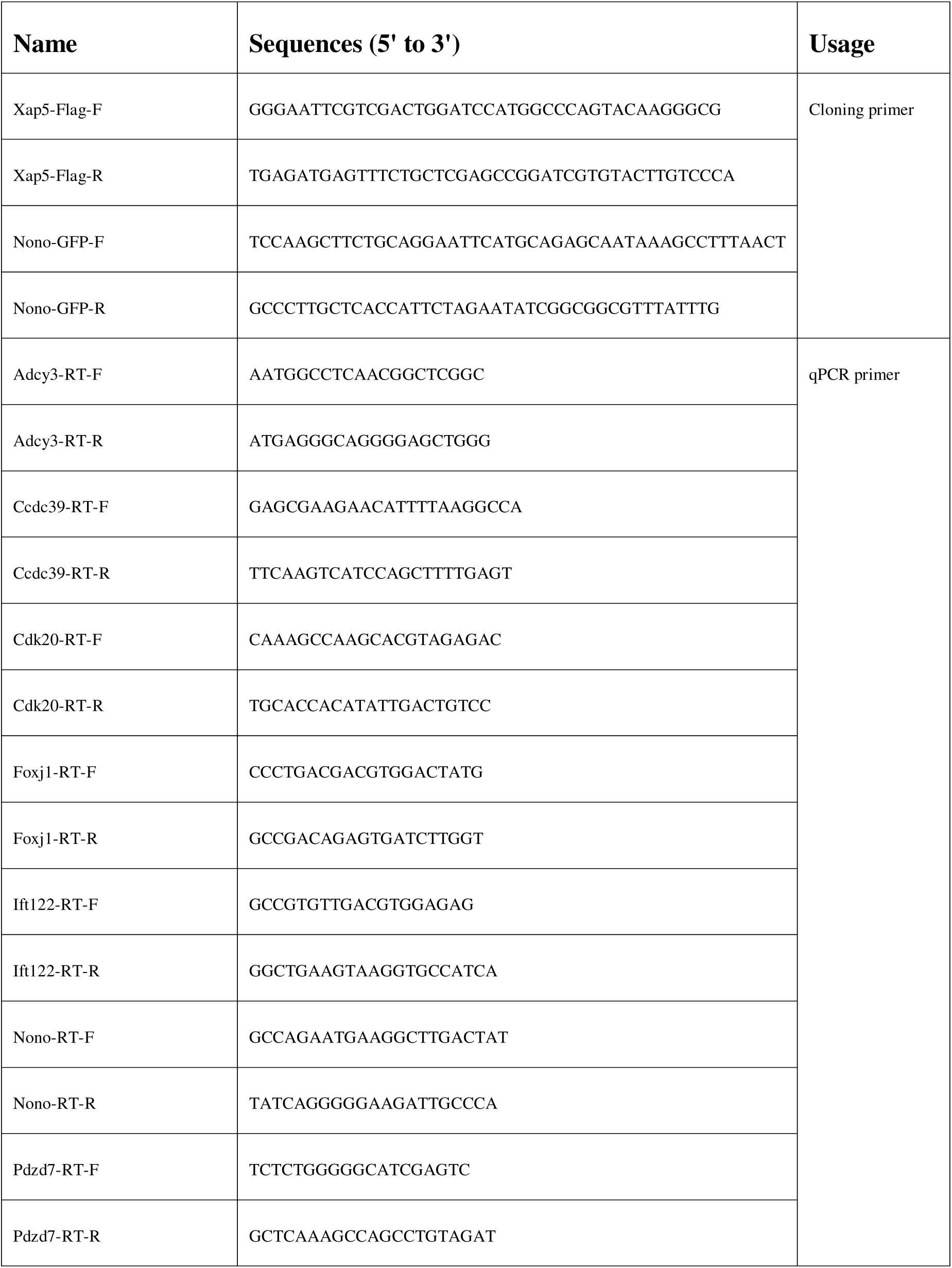
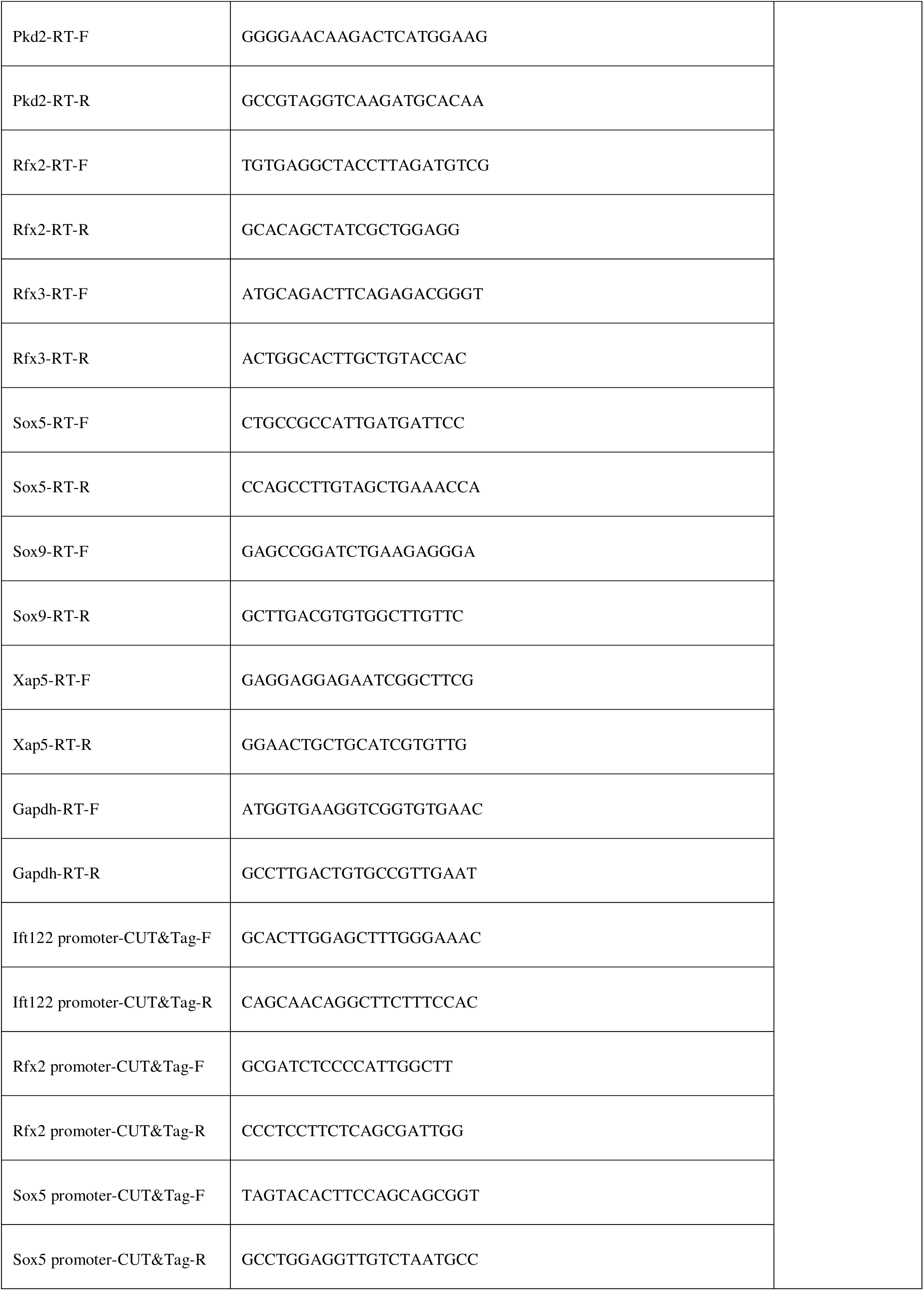
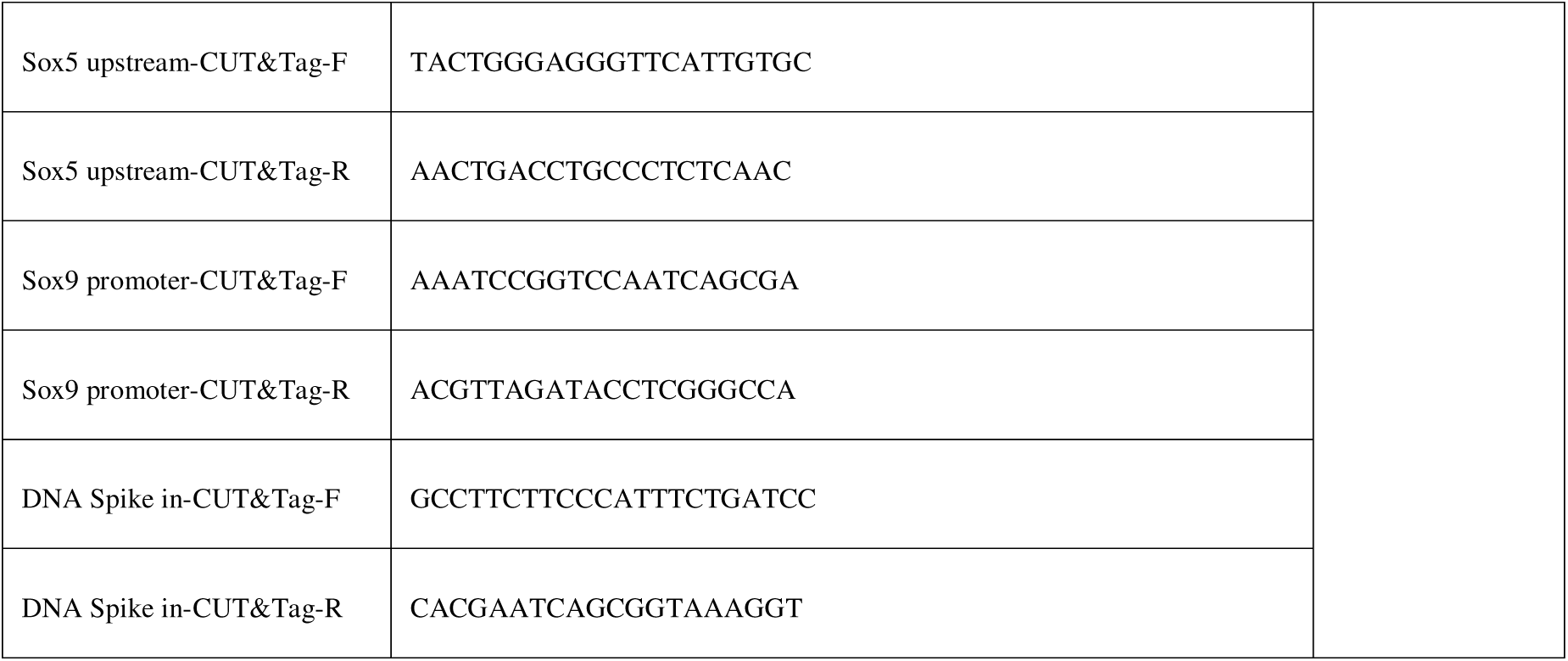
Primers used in the study.

**Supplementary Table 2.**
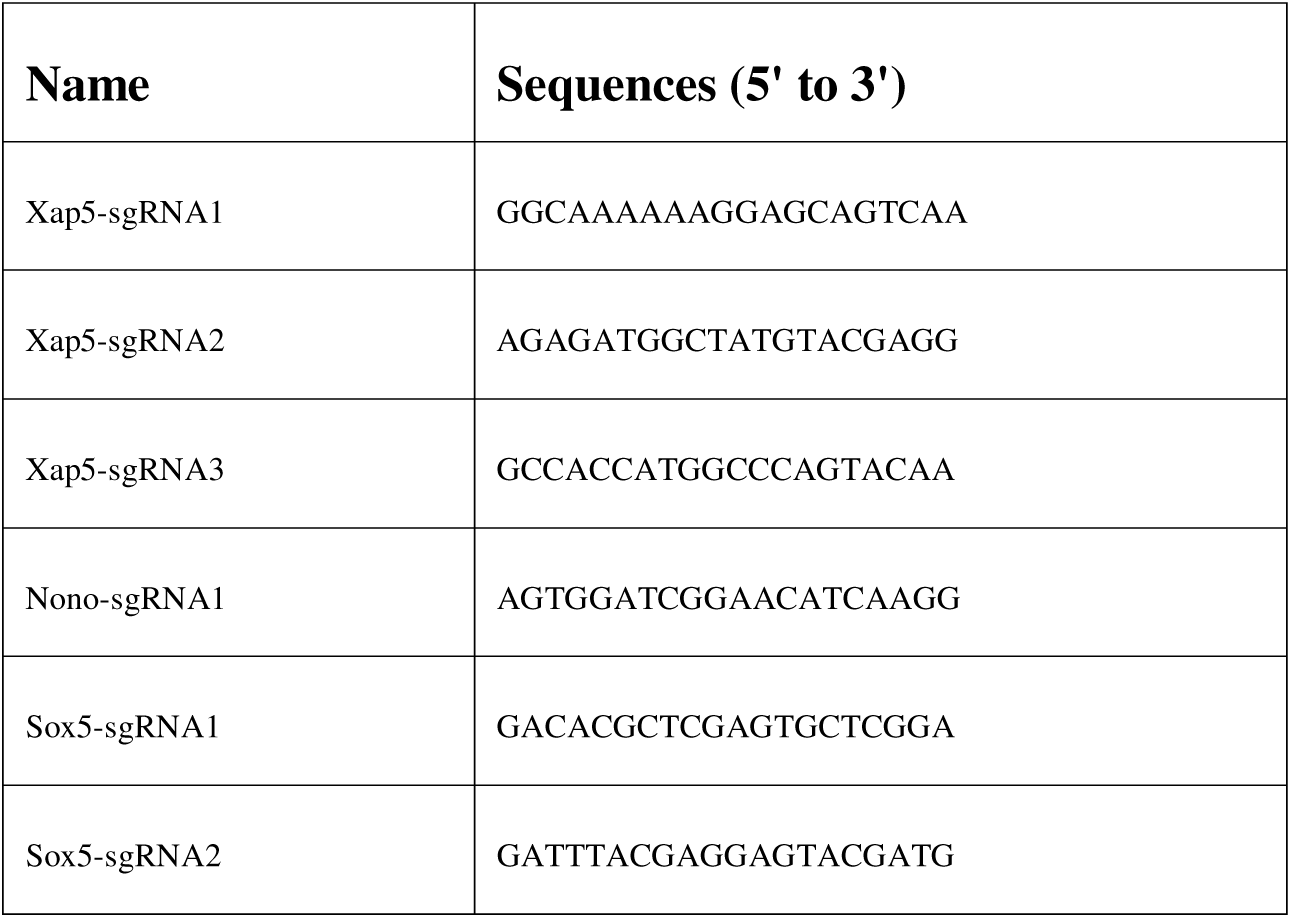
sgRNAs used in the study.

